# Neutrophil-derived extracellular vesicles: proinflammatory trails and anti-inflammatory microvesicles

**DOI:** 10.1101/583435

**Authors:** Young-Jin Youn, Sanjeeb Shrestha, Jun-Kyu Kim, Yu-Bin Lee, Jee Hyun Lee, Keun Hur, Nanda Maya Mali, Sung-Wook Nam, Sun-Hwa Kim, Dong-Keun Song, Hee Kyung Jin, Jae-sung Bae, Chang-Won Hong

## Abstract

Extracellular vesicles (EVs) are membrane-derived vesicles that mediate intercellular communications. Neutrophils produce different subtypes of EVs during inflammatory responses. Neutrophil-derived trails (NDTRs) are generated by neutrophils migrating toward inflammatory foci, whereas neutrophil-derived microvesicles (NDMVs) are thought to be generated by neutrophils that have arrived at the inflammatory foci. However, the physical and functional characteristics of neutrophil-derived EVs are incompletely understood. In this study, we investigated the similarities and differences between neutrophil-derived EV subtypes. Neutrophil-derived EVs shared similar characteristics regarding stimulators, generation mechanisms, and surface marker expression. Both neutrophil-derived EV subtypes exhibited similar functions, such as direct bactericidal activity and induction of monocyte chemotaxis via MCP-1. However, NDTR generation was dependent on the integrin signaling, while NDMV generation was dependent on the PI3K pathway. The CD16 expression level differentiated the neutrophil-derived EV subtypes. Interestingly, both subtypes showed different patterns of miRNA expression and were easily phagocytosed by monocytes. NDTRs induced M1 macrophage polarization, whereas NDMVs induced M2 macrophage polarization. Moreover, NDTRs but not NDMVs exerted protective effects against sepsis-induced lethality in a murine sepsis model and pathological changes in a murine chronic colitis model. These results suggest a new insight into neutrophil-derived EV subtypes: proinflammatory NDTRs and anti-inflammatory NDMVs.

**Key points:**

- Neutrophil-derived trails are proinflammatory extracellular vesicles that induce M1 macrophage polarization and protect against inflammation
- Neutrophil-derived microvesicles are anti-inflammatory extracellular vesicles that induce M2 macrophage polarization

## INTRODUCTION

Extracellular vesicles (EVs) are membrane-derived vesicles surrounded by lipid bilayers (Théry et al., 2009; >van der Pol et al., 2015). Since EVs express various ligands and receptors originating from their source cells and transport various molecules to their target cells (van der Pol et al., 2015), they are considered to be important mediators of intercellular communications. Neutrophils, the professional phagocytes, also produce EVs in response to various stimulators. Since the first discovery of membrane-containing vesicles released from neutrophils (Stein and Luzio, 1991), neutrophil-derived EVs have been called by different names, such as ectosomes (Duarte et al., 2012; Eken et al., 2010; 2013; Gasser et al., 2003; Hess et al., 1999), microparticles (Dalli et al., 2008; Gasser and Schifferli, 2005; Y. Hong et al., 2012; Kambas et al., 2014; Mesri and Altieri, 1999; 1998; Nieuwland et al., 2000; Nolan et al., 2008; Pitanga et al., 2014; Pliyev et al., 2014; Pluskota et al., 2008; Prakash et al., 2012; Slater et al., 2017), and microvesicles (Headland et al., 2015; Rhys et al., 2018; Timar et al., 2013). These vesicles are thought to be generated through membrane blebbing and shedding (Duarte et al., 2012; Nolan et al., 2008; Stein and Luzio, 1991) and conform to the general description of EVs (van der Pol et al., 2015). Recently, a specialized type of neutrophil-derived EVs was discovered and called trails (Lim et al., 2015). During migration through inflamed tissue, the uropods of neutrophils are elongated by adhesion to the endothelium (Hyun et al., 2012). These elongated uropods detach from the cell bodies, finally leaving chemokine-containing EVs (Lim et al., 2015). Hence, neutrophil-derived EVs can be categorized into two subtypes according to the mechanism of generation: neutrophil-derived microvesicles (NDMVs) and neutrophil-derived trails (NDTRs).

NMDVs are the classical type of neutrophil-derived EVs. Neutrophils generate NDMVs either spontaneously (Gasser and Schifferli, 2005; Mesri and Altieri, 1999) or in response to bacterial stimulation (Dalli et al., 2013; Eken et al., 2013; 2010; 2008; Hess et al., 1999; Mesri and Altieri, 1999; 1998; Pliyev et al., 2014; Pluskota et al., 2008; Slater et al., 2017; Timar et al., 2013). Various immunological stimuli, such as cytokines (Headland et al., 2015; Mesri and Altieri, 1999), chemokines (Pliyev et al., 2014), complement components (Pliyev et al., 2014; Stein and Luzio, 1991), and antibodies (Y. Hong et al., 2012; Kambas et al., 2014), also induce the production of NDMVs. Fluorescence-activated cell sorting (FACS) analysis and electron microscopic analysis showed that NDMVs are small (less than 1 μm in diameter) enclosed vesicles (Dalli et al., 2008; Duarte et al., 2012; Headland et al., 2015; Kambas et al., 2014; Pliyev et al., 2014; Prakash et al., 2012; Slater et al., 2017) surrounded with a double-layered membrane (Gasser et al., 2003; Hess et al., 1999). Phosphatidylserine is exposed in the outer layer of the membrane (Dalli et al., 2013; Eken et al., 2010; Gasser, 2004; Kambas et al., 2014; Nolan et al., 2008; Pitanga et al., 2014; Pliyev et al., 2014; Slater et al., 2017; Stein and Luzio, 1991; Timar et al., 2013) and various adhesion molecules, such as integrin αM (Dalli et al., 2013; Gasser et al., 2003; Y. Hong et al., 2012; Pluskota et al., 2008; Prakash et al., 2012; Slater et al., 2017), integrin β (Dalli et al., 2013), and L-selectins (Dalli et al., 2008; Gasser et al., 2003; Pitanga et al., 2014), are expressed on the surface. In addition, NDMVs contain protease-enriched granules (Dalli et al., 2013; Gasser et al., 2003; Hess et al., 1999; Y. Hong et al., 2012; Kambas et al., 2014; Mesri and Altieri, 1999; Pitanga et al., 2014; Slater et al., 2017) and express markers of neutrophil granules (Dalli et al., 2013; Gasser et al., 2003; Gasser and Schifferli, 2005; Headland et al., 2015; Hess et al., 1999; Y. Hong et al., 2012; Nieuwland et al., 2000; Pitanga et al., 2014; Prakash et al., 2012). The primary function of NDMVs is to modulate the inflammatory functions of neighboring cells. NDMVs enhance the expression of proinflammatory genes in endothelial cells (Y. Hong et al., 2012; Mesri and Altieri, 1998) and increase coagulation by inducing the aggregations of platelets and red blood cells (Gasser and Schifferli, 2005; Pluskota et al., 2008). NDMVs limit the growth of bacteria by inducing bacterial aggregation (Timar et al., 2013), deliver myeloperoxidase to endothelial cells (Mesri and Altieri, 1999), and increase phagocytosis by macrophages (Duarte et al., 2012). Moreover, NDMVs control inflammatory responses by modulating inflammatory gene expression in natural killer (NK) cells (Pliyev et al., 2014), monocyte-derived dendritic cells (moDCs) (Eken et al., 2008), macrophages (Eken et al., 2013; 2010; Gasser, 2004; Rhys et al., 2018), and chondrocytes (Headland et al., 2015). A recent study showed that NDMVs increased mortality in a murine model of sepsis by decreasing macrophage activation (Johnson et al., 2017).

NDTRs are a recently identified subtype of neutrophil-derived EV. In contrast to NDMVs, NDTRs are generated from neutrophils during extravasation from blood vessels into inflamed tissue (Hyun et al., 2012; Lim et al., 2015). Migrating neutrophils bind to endothelial cells via various adhesion molecules, including integrins (Hyun and C.-W. Hong, 2017; Nauseef and Borregaard, 2014). Therefore, physical forces between neutrophils and endothelial cells elongate the uropods of migrating neutrophils (Hyun et al., 2012), leading to the detachment of the tail portion (Lim et al., 2015). The detached tail portion contains CXC-chemokine ligand 12 (CXCL12) which guides CD8^+^ T cells to the influenza-infected tissues (Lim et al., 2015).

Despite these advances in our understanding of neutrophil-derived EVs, the similarities and differences between NDTRs and NDMVs are not fully understood. The most important difference between NDTRs and NDMVs seems to be the spatiotemporal generation mechanism; NDTRs are generated from neutrophils migrating toward inflammatory foci, whereas NDMVs are generated from neutrophils that have arrived at inflammatory foci. Based on this difference, we hypothesized that NDTRs and NDMVs play different roles in modulating immune responses. Here, we investigated the similarities and differences between NDTRs and NDMVs. Although neutrophil-derived EVs were indistinguishable with respect to several characteristics, we found that NDTRs exert proinflammatory effects, whereas NDMVs exert anti-inflammatory effects.

## RESULTS

### Various stimulators induce the formation of both NDTRs and NDMVs

To identify specific stimulators that induce the formation of NDTRs and NDMVs, we examined the effects of various stimulators on formation of these EVs. The stimulators were categorized into the following subgroups: (i) pathogen-associated molecular patterns (PAMPs): formyl-methionyl-leucyl-phenylalanine (fMLP), lipopolysaccharide (LPS), opsonized *E. coli*, and opsonized *S. aureus*; (ii) danger-associated molecular patterns (DAMPs): high mobility group box 1 (HMGB1), complement component 5a (C5a), and S100 calcium binding protein B (S100B); (iii) inflammatory cytokines: tumor necrosis factor-α (TNF-α) and interferon-γ (IFN-γ); (iv) anti-inflammatory or immunosuppressive cytokines: tumor growth factor-β (TGF-β) and interleukin-4 (IL-4); and (v) exogenous compounds: N(γ)-nitro-L-arginine methyl ester (L-NAME) and phorbol 12-myristate 13-acetate (PMA). To visualize EV formation, neutrophils were stained with cell tracker green and live-cell fluorescence images were obtained. All stimulators induced the formation of NDTRs from neutrophils in the fibronectin-coated chemotaxis chamber (Figure 1A, Supplementary movie 1). The stimulated neutrophils exhibited elongated uropods (Figure 1A, arrows), which were subsequently detached from the cell bodies, forming NDTRs (Figure 1A, arrowheads). Intriguingly, all stimulators also induced the formation of NDMVs from neutrophils in the uncoated confocal dish (Figure 1A, Supplementary movie 2). Live-cell fluorescence imaging showed the membrane blebbing and shedding of membranes (Figure 1A, arrowheads), the membranes finally detached, forming NDMVs. The relative amounts of NDTRs and NDMVs were quantified using a spectrofluorometer (Figure 1B and 1C). These results suggest that the generation of NDTRs and NDMVs by the abovementioned stimulators is indistinguishable.

**Figure 1.**
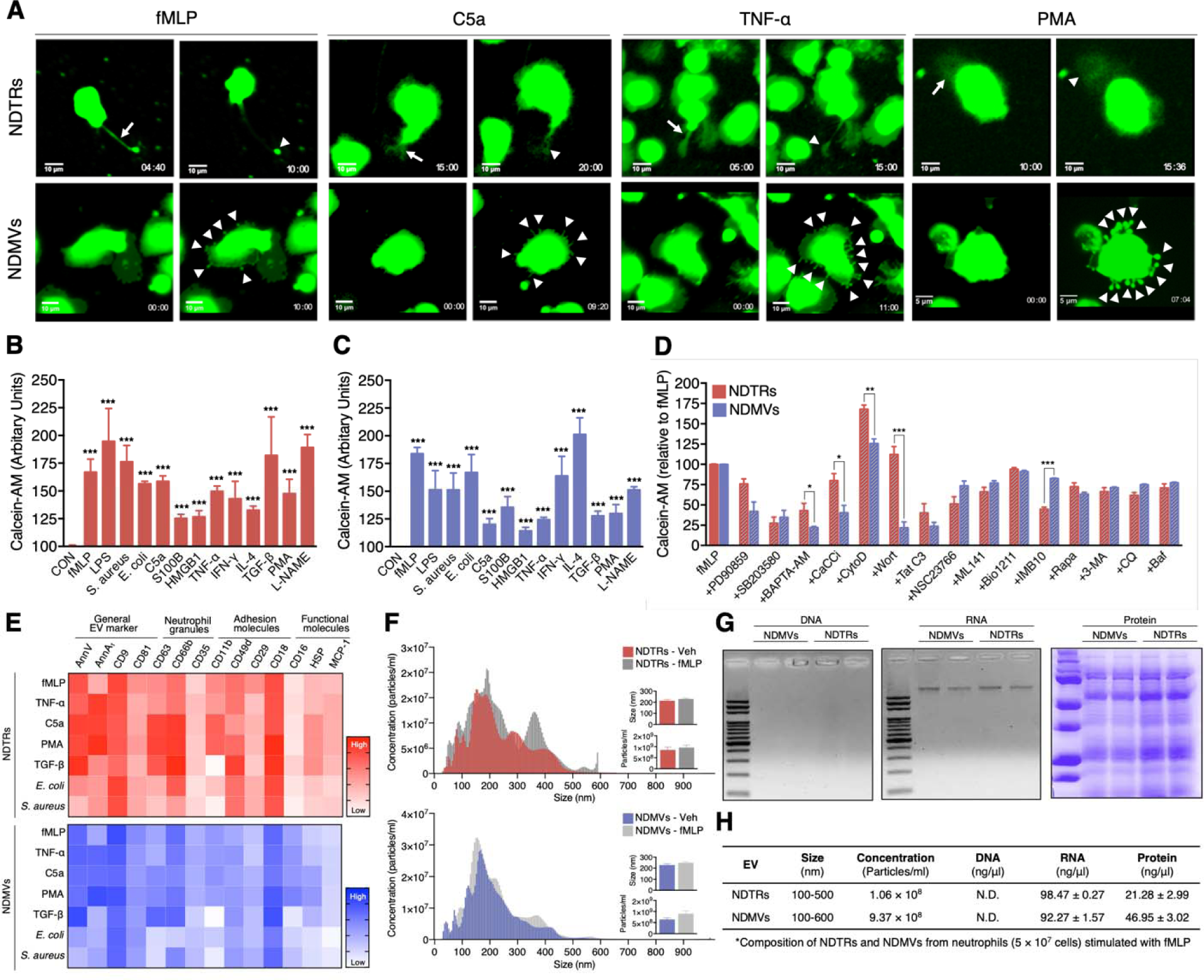
Characterization of NDTRs and NDMVs. (A) Representative time-lapse images of EVs released from neutrophils. The time denotes minutes after stimulation. The arrows indicate elongated uropods. The arrowheads indicate deposited neutrophil-derived EVs. (B-D) Quantification of relative amounts of generated neutrophil-derived EVs. Neutrophils were stained with calcein-AM and stimulated with the indicated stimulators. n = 4–7 per group. (B-C) Relative fluorescence normalized to the fluorescence of EVs isolated from unstimulated neutrophils. (D) Neutrophils were stimulated with fMLP in the presence or absence of th indicated inhibitors. Relative fluorescence normalized to the fluorescence of EVs isolated from fMLP-stimulated neutrophils. (E) Heatmap of marker expression levels. n = 3. (F) Representative density plots of neutrophil-derived EVs. (G-H) Validation of the contents in neutrophil-derived EVs. All data are representative of more than three independent experiments with n = 3 per group. The data shown are the mean ± SEM. (B-C) ***P < 0.001 vs control. (D) *P < 0.05; **P < 0.01; ***P < 0.001.

To further investigate the mechanism underlying NDTR and NDMV formation, we examined the effects of various inhibitors on the formation these EVs (Figure 1D). Neutrophils were allowed to generate neutrophil-derived EVs in response to fMLP in the presence or the absence of various signaling pathway inhibitors. PD90859 (an ERK inhibitor), SB203580 (a p38 MAPK inhibitor), BAPTA-AM (a calcium chelator), CaCCinh (a Ca^2+^-activated Cl^-^ channel transmembrane protein 16A inhibitor), Tat-C3 (a Rho inhibitor), NSC23766 (a Rac1 inhibitor), and ML141 (a Cdc42 inhibitor) attenuated the formation of both NDTRs and NDMVs (Figure 1E). However, wortmannin [a phosphatidylinositol 3-kinase (PI3K) inhibitor] significantly attenuated the formation of NDMVs but not NDTRs (Figure 1D). Moreover, IMB10 (an inhibitor of integrin αM and MAC-1) significantly inhibited the formation of NDTRs but slightly attenuated the formation of NDMVs (Figure 1D). These results suggest that NDTRs and NDMVs share a common mechanism of generation but that integrin signaling and PI3K-dependent exocytosis pathways are required for the generation of NDTRs and NDMVs, respectively.

### CD16 expression differentiates NDTRs from NDMVs

Next, we assessed the expression of surface markers on NDMVs and NDTRs. FACS analysis indicated that both NDTRs and NDMVs expressed the surface markers of general EVs (Lötvall et al., 2014), such as annexins [Annexin V (AnnV) and annexin A1 (AnnA_1_)] and tetraspanins (CD9 and CD81) in response to various stimulators (Figure 1E, Supplementary Figure 1). To further confirm that these EVs were neutrophil-specific, the expression of neutrophil-specific markers on EVs isolated from fMLP-stimulated neutrophils was examined. Both NDTRs and NDMVs isolated from fMLP-stimulated neutrophils expressed high levels of neutrophil granule markers: CD63 (primary granules), CD66b (secondary granules), and CD35 (tertiary granules) (Figure 1E). The adhesion molecule expression analysis showed that all integrin markers were expressed at high levels in neutrophil-derived EVs: CD11b (integrin αM), CD49d (integrin α4), and CD29 (integrin β1), and CD18 (integrin β2) (Figure 1E). Moreover, both NDTRs and NDMVs contained heat shock protein (HSP70) and monocyte chemoattractant protein 1 (MCP-1) (Figure 1E). Interestingly, NDTRs and NDMVs showed different levels of CD16 (an Fcγ type III receptor) expression. CD16 expression was high in NDMVs but negligible in NDTRs (Figure 1E). Neutrophil-derived EVs isolated from neutrophils stimulated with C5a, TNF-α, or PMA showed surface marker expression patterns similar to those of EVs isolated from fMLP-simulated neutrophils (Figure 1E). Notably, the CD16 expression level was consistently higher in NDMVs than in NDTRs. Bacteria-induced NDTRs and NDMVs showed the similar patterns but reduced extents of surface marker expression. TGF-β-induced NDTRs and NDMVs showed similar patterns of surface marker expression except for the reciprocal expression of CD16 in NDTRs (Figure 1E).

Next, we evaluated the sizes and numbers of NDTRs and NDMVs. Nanoparticle tracking analysis (NTA) revealed heterogeneity in the size distribution of spontaneously generated neutrophil-derived EVs (Figure 1F). The diameter of spontaneously generated neutrophil-derived EVs was between 50 and 600 nm, with a mean diameter of 200 nm. The sizes and diameters of neutrophil-derived EVs isolated from fMLP-simulated neutrophils were not significantly different from those of spontaneously generated neutrophil-derived EVs (spontaneous vs fMLP-simulated: NDTR, 246.7 ± 10.80 nm vs 226.5 ± 14.11 nm; NDMV, 227.2 ± 9.32 nm vs 211.5 ± 13.47 nm). To further identify the characteristics of NDTRs and NDMVs, we investigated their contents and fount that both subtypes of EVs contained RNA and protein but not DNA (Figure 1G and Supplementary Figure 2A). The concentrations of proteins retained in NDTRs and NDMVs varied according to the stimulation (Supplementary Figure 2B and 2C). Collectively, these results suggest that neutrophil-derived EVs are defined as AnnV^+^ AnnA1^+^ CD9^+^ CD63^+^ CD66b^+^ CD35^+^ CD11b^+^ CD49d^+^ CD29^+^ CD18^+^ EVs that are loaded with RNA and proteins. Furthermore, the expression levels of CD16 differentiated NDTRs (CD16^low^) from NDMVs (CD16^High^).

**Figure 2.**
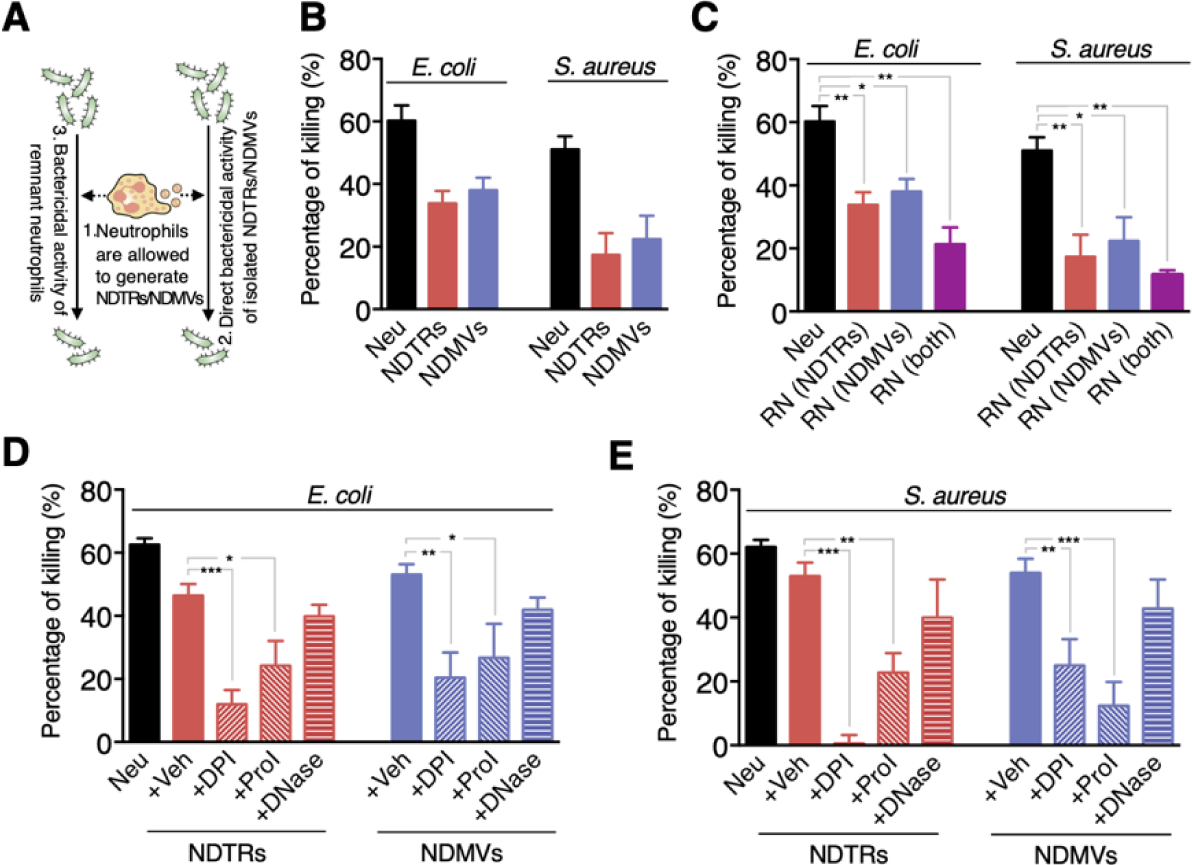
NDTRs and NDMVs exert direct bactericidal activity. (A) Schematic of the experiments. (B) Direct bactericidal activity of neutrophil-derived EVs. (C) Bactericidal activity of RNs after the formation of neutrophil-derived EVs. (D-E) The effects of inhibitors of the specific bactericidal activity pathways on the bactericidal activity of neutrophil-derived EVs. The data are pooled from three independent experiments (n = 4–7 per each group) and shown as the mean ± SEM. *P < 0.05; **P < 0.01; ***P < 0.001.

### Both NDTRs and NDMVs have direct bactericidal activity

A previous study reported that NDMVs inhibit the growth of bacteria by inducing bacterial aggregation (Timar et al., 2013). Since neutrophil-derived EVs expressed neutrophil granules, we next examined the bactericidal activity of these EVs. Neutrophils were stimulated with either *E. coli* or *S. aureus*, and EVs were isolated. Then, *E. coli* and *S. aureus* were exposed to equivalent amounts of neutrophil-derived EVs, and the direct bactericidal activity of the neutrophil-derived EVs was assessed (Figure 2A). Both NDTRs and NDMVs showed significant bactericidal activity against *E. coli* and *S. aureus* (Figure 2B). Neutrophil-derived EVs are composed of plasma membranes and carry substantial amounts of proteins and RNA, hence, we hypothesized that the continual production of neutrophil-derived EVs might negatively influence the overall bactericidal efficiency of remnant neutrophils (RNs). Therefore, we examined the bactericidal activity of RNs after the generation of neutrophil-derived EVs. Neutrophils were allowed to generate either NDTRs, NDMVs, or both, and the bactericidal activity of RNs was examined (Figure 2A). Interestingly, RNs exhibited the significantly diminished bactericidal activity against *E. coli* and *S. aureus* (Figure 2C), suggesting that neutrophil-derived EVs might be involved in the general bactericidal activity of neutrophils.

Neutrophils exert their bactericidal activity through the release of reactive oxygen species (ROS), granules, and neutrophil extracellular traps (NETs) (Nauseef and Borregaard, 2014). We thus evaluated the mechanism underlying the bactericidal activity of neutrophil-derived EVs by inhibiting each pathway of bactericidal activity. The NADPH oxidase inhibitor (diphenyleneiodonium, DPI) and protease inhibitor (PI) cocktail significantly attenuated the bactericidal activity of neutrophil-derived EVs (Figure 2D and 2E). However, DNase, an inhibitor of NETs, did not inhibit the bactericidal activity of neutrophil-derived EVs (Figure 2D and 2E). A luminol assay showed that both NDTRs and NDMVs generate ROS in response to PMA stimulation (Supplementary Figure 2E). Collectively, these results suggest that both NDTRs and NDMVs exert bactericidal activity via ROS- and granule-dependent mechanisms.

### Both NDMVs and NDTRs induce monocyte chemotaxis

The primary function of NDTRs is to guide influenza-specific CD8^+^ T cells to the site of infection through retained CXCL12 (Lim et al., 2015). Since we found MCP-1 expression in NDTRs and NDMVs, the effects of neutrophil-derived EVs on the chemotaxis of monocytes and macrophages were examined. One side of a chemotaxis chamber was coated with NDTRs by allowing neutrophils to move toward various chemoattractants (Figure 3A, left panel). Monocytes and macrophages were isolated from peripheral mononuclear cells (PBMCs), and their chemotaxis toward NDTRs was examined (Figure 3B, direct NDTR). Monocytes showed significant chemotaxis in NDTR-coated chemotaxis chamber (Figure 3B, direct NDTR), while macrophages did not exhibit significant movements (Supplementary Figure 3A and 3B). Moreover, in a chemotaxis chamber coated with isolated NDTRs and NDMVs, monocytes exhibited efficient chemotaxis toward neutrophil-derived EVs (Figure 3A and 3B, isolated NDTRs/NDMVs). However, the pharmacological inhibition of CXCR2, a receptor for MCP-1, completely abolished the migration of monocytes toward to NDTRs and NDMVs (Figure 3B, +MCP-1 inhibitor). These results suggest that neutrophil-derived EVs induce monocyte chemotaxis in an MCP-1-dependent manner. Additionally, NDTRs and NDMVs induce neutrophil chemotaxis and LY223982 (a leukotriene B_4_ antagonist) completely inhibited neutrophil chemotaxis (Supplementary Figure 3C and 3D).

**Figure 3.**
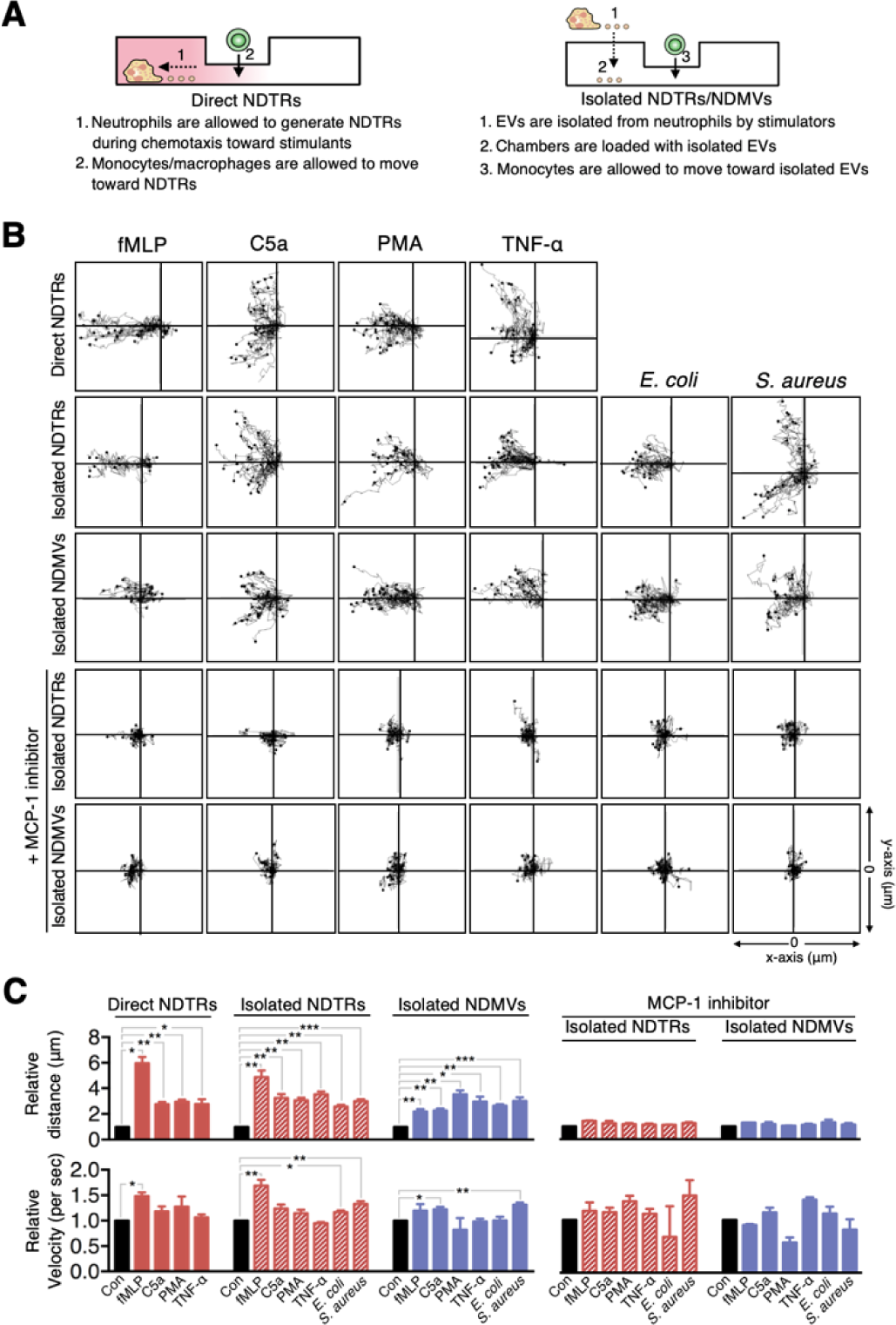
NDTRs and NDMVs induce monocyte chemotaxis via an MCP-1-dependent pathway. (A) Schematic of the chemotaxis experiments. (B) Monocyte migration tracking analysis. The distances traveled by migrating cells were tracked on every minute for 45 min. Representative tracking results of thirty cells per each group are presented. (C) Relative mean distance and relative mean velocity of monocytes traveled towards neutrophil-derived EVs. (B-C) Direct NDTRs: neutrophils were allowed to generate NDTRs in a fibronectin-coated μ-slide chamber, and the movements of monocytes was measured. Isolated NDTRs and NDMVs: the lane was coated with isolated neutrophil-derived EVs, and the movement of monocytes wa measured. MCP-1 inhibition: monocytes were loaded into the lane in the presence of a CCR-2 antagonist. The data are shown as the mean ± SEM. *P < 0.05; **P < 0.01; ***P < 0.001. All data are representative of three independent experiments (n = 3–4 per group).

### NDTRs and NDMVs differentially induce the polarization of monocytes into different phenotypes

Previous studies have shown that NDMVs exert anti-inflammatory responses by suppressing the activation of immune cells such as dendritic cells (Eken et al., 2008), monocytes (Prakash et al., 2012), and macrophages (Duarte et al., 2012; Eken et al., 2013; 2010; Gasser, 2004; Johnson et al., 2017). Since NDMVs are generated from neutrophils in tissues (Eken et al., 2013; Headland et al., 2015; Rhys et al., 2018; Slater et al., 2017), these EVs might be involved in limiting excessive inflammation by suppressing neighboring immune cells. However, NDTRs are produced during extravasation toward inflammatory foci (Hyun et al., 2012; Lim et al., 2015); hence, we hypothesized that NDTRs might stimulate the following immune cells to prepare for pathogen defense. Therefore, we compared the effects of NDTRs and NDMVs on the phenotypic polarization of monocytes into macrophages.

To evaluate this hypothesis, we first investigated the direct interactions between neutrophil-derived EVs and monocytes. THP-1 cells, human monocytic cell line, were differentiated into M0 macrophages by stimulation with PMA (100 ng/ml, 48 h) and were exposed to neutrophil-derived EVs in a microfluidic chamber. M0 cells actively migrated toward neutrophil-derived EVs and phagocytosed them (Figure 4A, Supplementary movie 3 and 4). The phagocytosis of neutrophil EVs by M0 macrophages was further confirmed by FACS analysis (Figure 4B).

**Figure 4.**
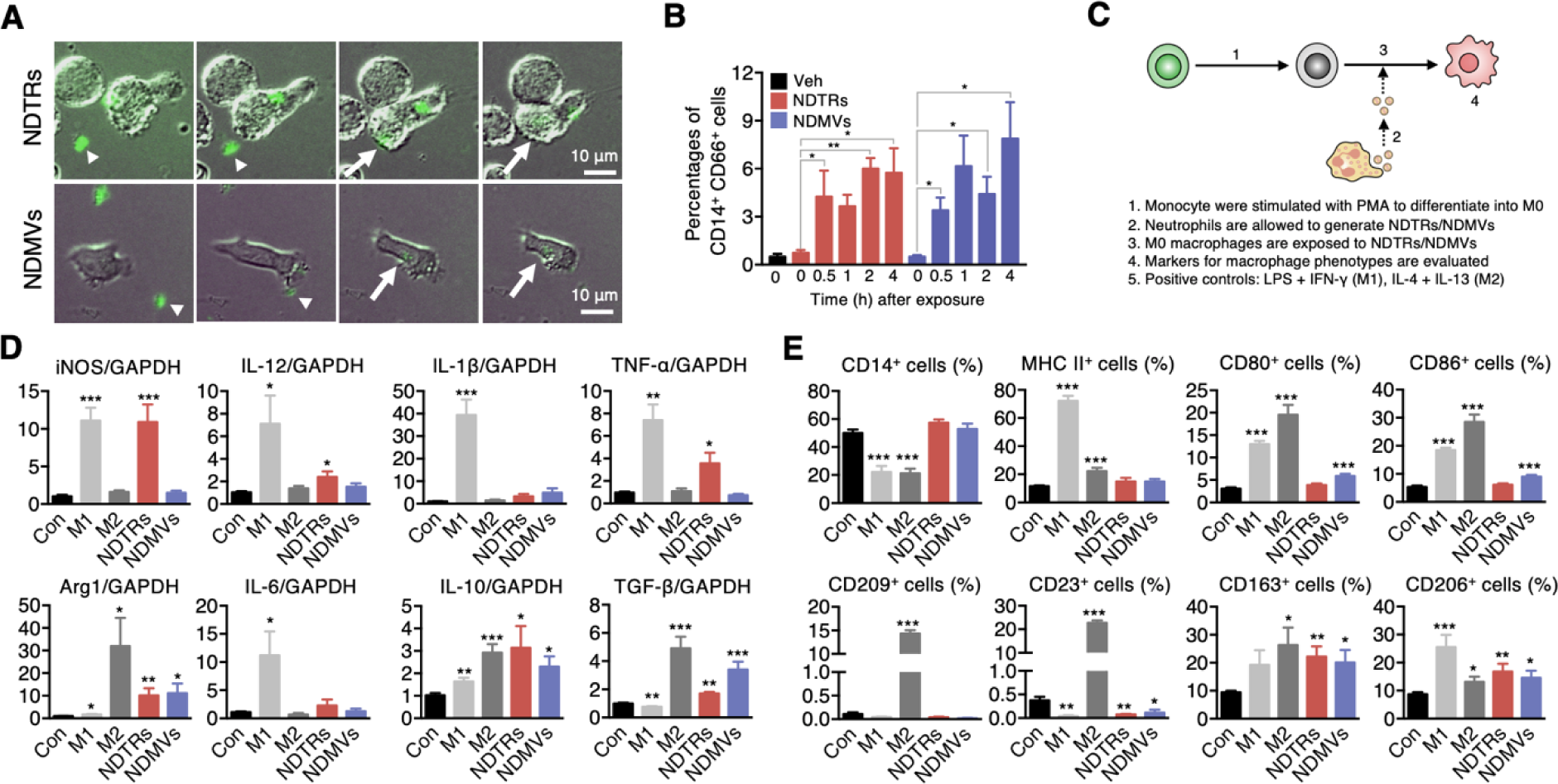
Differential effects of NDTRs and NDMVs on macrophage phenotyp polarization. (A) Time-lapse images of M0-differentiated THP-1 cells acquiring neutrophil-derived EVs. Green, neutrophil-derived EVs stained with Cell Tracker Green. Arrowheads, neutrophil-derived EVs attached to the plates. Arrows, neutrophil-derived EVs phagocytosed by M0-differentiated THP-1 cells. Representative images of three independent experiments. (B) FACS analysis showing the uptake of neutrophil-derived EVs by M0-differentiated THP-1 cells. n = 3. (C) Schematic of the phenotype polarization experiments. (D-E) The expression of M1 and M2 macrophage-associated markers in M0-differentiated THP-1 cells exposed to neutrophil-derived EVs (n = 3–4 per each group). The data are shown as the mean ± SEM. *P < 0.05; **P < 0.01; ***P < 0.001 compared to control.

Next, we evaluated the effects of NDTRs and NDMVs on the phenotypic polarization of macrophages. THP-1 cells were differentiated into M0 macrophages and further exposed to either NDTRs or NDMVs (Figure 4C). For comparison, M0-differentiated THP-1 cells were incubated with either LPS/IFN-γ or IL-4/IL-13 for differentiation into M1 or M2 macrophages, respectively (Figure 4C). Interestingly, NDTRs and NDMVs differentially induced the phenotypic polarization of M0-differentiated THP-1 cells. NDTR-exposed M0 cells showed increased expression of phenotypic markers of proinflammatory macrophages, such as inducible nitric oxide synthase (iNOS), IL-12, and TNF-α (Figure 4D and 4E). However, NDTR exposure also induced the upregulation of phenotype markers of anti-inflammatory macrophages, such as arginase-1 (Arg-1), IL-10 and TGF-β (Figure 4D and 4E). Furthermore, NDTR-treated M0 macrophages showed the reduced expression of CD23 with increased expression of CD163 and CD206 (Figure 4D and 4E). Although these results indicate that NDTR-treated macrophages did not exhibit the classical patterns of M1/M2 markers, recent studies suggest that the surface marker expression patterns of macrophages polarized toward M1 by exogenous compounds do not always follow those for classical M1 macrophages (Lee et al., 2017; Xu et al., 2016; Zhang et al., 2018). Therefore, we defined the phenotype of NDTR-treated macrophages as M1-like macrophages based on the expression of important functional markers of M1 macrophages, such as iNOS, TNF-α, and IL-12, in NDTR-treated macrophages.

However, NDMV-exposed M0 macrophages showed different patterns of phenotypic marker expression. NDMV-treated M0 macrophages showed the increased expression of anti-inflammatory markers such as TGF-β, IL-10, and Arg-1, with increased expression of CD163, CD206, CD80, and CD86 (Figure 4D and 4E). Moreover, these cells showed reduced expression of CD23 (Figure 4E) and lacked the expression of CD1a (data not shown). Collectively, these results suggest that NDMVs induce the differentiation of M0 macrophages into M2 macrophages, which are important for tissue remodeling and immunoregulation (Martinez and Gordon, 2014). When undifferentiated THP-1 cells were directly exposed to neutrophil-derived EVs, neither their morphology (Supplementary Figure 4B) nor their expression of M1/M2 markers (Supplementary Figure 4C-D) significantly changed.

### miRNAs are differentially expressed in NDTRs and NDMVs

miRNAs are small noncoding RNA molecules that posttranscriptionally regulate gene expression. Various miRNAs are found in EVs and are considered to be important mediators of intercellular communications (Camussi et al., 2010; Valadi et al., 2007; van der Pol et al., 2012). Since recent studies indicate that miRNAs play a pivotal role in the phenotypic polarization of macrophages (Graff et al., 2012; O’Connell et al., 2010; Saha et al., 2016), we performed miRNA sequencing (miRNA-seq) on neutrophil-derived EVs. Agilent Human miRNA Microarray assay was performed on the neutrophil-derived EVs isolated from six volunteers (Figure 5A). The hierarchical clustering shows the differential miRNA expression patterns between NDTRs and NDMVs (Figure 5B). Furthermore, analysis of differentially expressed miRNAs (adjusted p-value < 0.05, fold change ≥ 1.5) identified 40 miRNAs with significantly different expression levels in NDTRs and NDMVs (Figure 5A). Among the top 10 differentially expressed miRNAs, six miRNAs (miR-122-5p, miR-1260a, miR-1285-5p, miR-24-3p, miR-29a-3p, and miR-4454+miR7975) were highly expressed in NDTRs, while three miRNAs (miR-126-3p, miR-150-5p, and miR-451a) were highly expressed in NDMVs (Figure 5C). Next, qRT-PCR analysis was conducted to validate the differential miRNA expression in neutrophil-derived EVs. Cel-miR-39 was added to the samples during the RNA extraction step and was used as an exogenous control for normalization. We found that four miRNAs (miR-1260, miR-1285, miR-4454, and miR-7975) were highly expressed in NDTRs, while three miRNAs (miR-126, miR-150, and miR-451a) were highly expressed in NDMVs (Figure 5D). To further confirm the involvement of miRNAs in macrophage polarization, M0-differentiated THP-1 cells were transfected with mimics of miRNAs differently expressed in NDTRs and NDMVs. Interestingly, the transfection with miRNA mimics highly expressed in NDTRs (miR-1260a, miR-1285-5p, miR-4454, and miR-7975) enhanced iNOS expressions in M0-differentiated THP-1 cells (Figure 5E). In contrast, the transfection with miRNA mimics highly expressed in NDMVs (miR-150-5p and miR-451a) enhanced arginase expressions in M0-differentiated THP-1 cells (Figure 5E).

**Figure 5.**
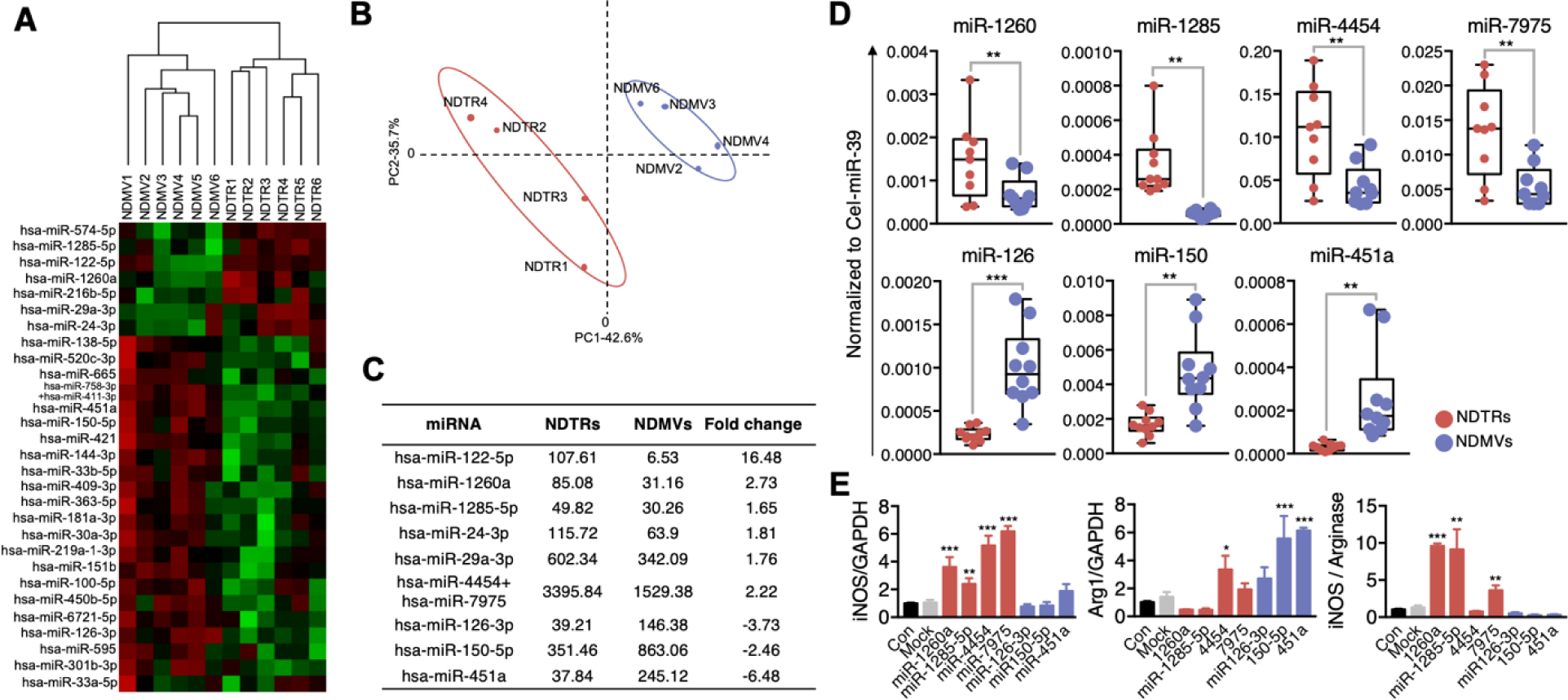
Differential miRNA profiles in NDTRs and NDMVs. (A) Hierarchical clustering of differentially expressed miRNAs in NDTRs and NDMVs. The miRNA profiles of NDTRs and NDMVs (n = 6 per group) clustered. Cluster analysis based on log10-transformed data. A red color represents relatively higher expression and a green color represents relatively lower expression. (B) Principal component analysis (PCA) plot of the miRNA expression profiles of NDTRs and NDMVs. (C) Summary of selected miRNAs differentially expressed in NDTRs and NDMVs. (D) Validation of selected miRNAs in NDTRs and NDMVs using RT-PCR (n = 10 per each group). (E) The effects of miRNA mimics (miR-1260a, miR-1285-5p, miR-4454, miR-7975, miR-126-3p, miR-150p, and miR-451a) on the iNOS and arginase expression in M0-differentiated THP-1 cells (n = 3). The data are shown as the mean ± SEM. *P < 0.05; **P < 0.01; ***P < 0.001.

### NDTRs exert protective effects against acute and chronic inflammation

Because NDTRs and NDMVs exerted different effects on the phenotypic polarization of macrophages, we next evaluated the clinical implication of NDTRs and NDMVs in the murine models of acute and chronic inflammation. We employed an experimental sepsis model (cecal-ligation and puncture, CLP) to establish murine model of acute inflammation (Park et al., 2017) and a chronic dextran sulfate sodium (DSS)-induced colitis model to establish a murine model of chronic inflammation (Marcon et al., 2013). BALB/c mice were treated intraperitoneally with either NDTRs or NDMVs 30 min prior to CLP surgery and further treated on days 1, 2, and 3 after CLP surgery. NDTR-treated mice showed increased survival, while NDMV-treated mice did not show any survival benefit (Figure 6A). To establish a chronic inflammation model, mice were orally administered with two cycles of 2% DSS as previously described (Marcon et al., 2013). Either NDTRs or NDMVs were administered intraperitoneally on days 16, 18, 20, and 22 (Figure 6B). Interestingly, NDTRs significantly reduced colon length, while NDMVs had no effect (Figure 6C). Moreover, NDTR treatment significantly reduced colon damage (Figure 6D).

**Figure 6.**
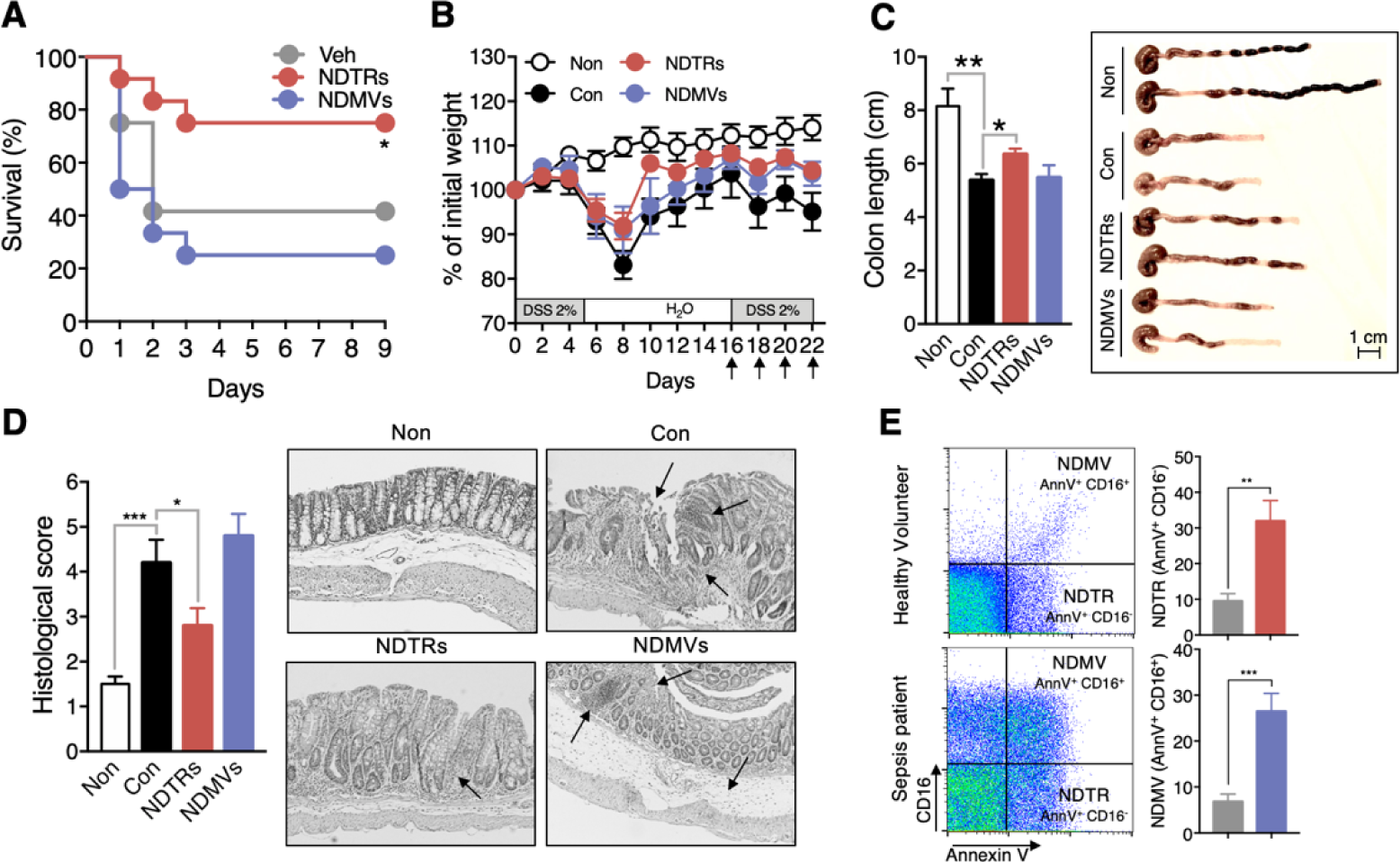
Differential effects of NDTRs and NDMVs in murine models of inflammation. (A) Effects of NDTRs and NDMVs in a murine model of sepsis. Experimental sepsis was induced by CLP. (A) Survival rates of septic mice after the administration of neutrophil-derived EVs. BALB/c mice were administered an i.p. injection of either NDTRs (n = 8), NDMVs (n = 12) or vehicle (saline, n = 12) 1 h before surgery and on days 1 and 3 after surgery. *P < 0.05 compared to vehicle. (B-E) The effects of NDTRs and NDMVs in a murine model of chronic colitis. Mice were administered with two cycles of 2% DSS for 5 days. At the beginning of the second cycle, mice were treated with NDTRs and NDMVs on every 2 days. n = 4 per each group. (B) Percentage of initial weight. (C-D) Mice were sacrificed on day 22 and subjected to evaluation. (C) Left panel, colon length. Right panel, representative photographs of the colon and cecum. (D) Macroscopic colonic damage. Left panel, histological score. Right panel, representative image of hematoxylin and eosin (H&E) staining. The arrows indicate crypt damage and inflammatory cell infiltration. (E) The identification of neutrophil-derived EVs in plasma of sepsis patients. EVs were isolated from plasma of either healthy volunteers (n = 4) or sepsis patients (n = 12), and stained with AnnV and CD16. Left panel, representative scatter plot. Right panel, the percentages of NDTRs and NDMVs.

To further confirm the clinical relevance of neutrophil-derived EVs, we examined the existence of NDTRs and NDMVs in serum of sepsis patients. Serum was obtained from 12 septic patients admitted to the intensive care units due to community-acquired pneumonia. Interestingly, the percentages of both NDTRs (AnnV^+^ CD16^-^) and NDMVs (AnnV^+^ CD16^+^) were higher in serum of sepsis patients than in that of healthy volunteers (Figure 6E).

## DISCUSSION

In this study, we investigated the similarities and differences between neutrophil-derived EVs and found that NDTRs and NDMVs have many similarities. Both were generated by the same stimulators (Figure 1A-C), shared the mechanism of generation (Figure 1D), and showed similar marker expression patterns (Figure 1E) and physical characteristics (Figure 1F-1H). Moreover, both NDTRs and NDMVs exerted similar functions. Both showed direct bactericidal activity (Figure 2), induced monocyte chemotaxis via MCP-1 (Figure 3) and were phagocytosed by monocytes (Figure 4A and 4B). However, we found profound differences between NDTRs and NDMVs. First, NDMVs were distinguished from NDTRs by the expression levels of CD16 (Figure 1E). In addition, although NDTRs and NDMVs share a common mechanism of generation, the generation of NDTRs depends on integrin signaling, while the generation of NDMVs depends on the PI3K pathway (Figure 1D). Moreover, NDTRs and NDMVs showed different patterns of miRNA expression (Figure 5A-E) and induced the phenotypic polarization of monocytes into either proinflammatory or anti-inflammatory macrophages, respectively (Figure 4C-E, 5E). Finally, only NDTRs showed beneficial effects in murine models of acute and chronic inflammation (Figure 6).

“EVs” is an umbrella term encompassing all types of vesicles derived from prokaryotic and eukaryotic cells (van der Pol et al., 2015); hence, EVs consist of diverse subpopulations that can be discriminated by sizes, contents, and functions (Kowal et al., 2016; Tkach et al., 2017). Since NDTRs were discovered only recently, the detailed rationale for the classification of NDTRs as a distinct subtype of neutrophil-derived EVs is mandatory. Therefore, we first sought specific stimulators of NDTR and NDMV production. Previous studies identified various stimulators of NDMV generation: fMLP (Dalli et al., 2013; Eken et al., 2013; 2010; 2008; Hess et al., 1999; Mesri and Altieri, 1999; 1998; Pliyev et al., 2014), LPS (Pluskota et al., 2008), C5a (Pliyev et al., 2014; Stein and Luzio, 1991), TNF-α (Headland et al., 2015; Timar et al., 2013), IL-8 (Headland et al., 2015; Mesri and Altieri, 1999; Pliyev et al., 2014), antinuclear cytoplasmic antibody (Y. Hong et al., 2012; Kambas et al., 2014), PMA (Headland et al., 2015; Hess et al., 1999; Mesri and Altieri, 1999; Pluskota et al., 2008; Slater et al., 2017; Timar et al., 2013), zymosan (Pliyev et al., 2014), mycobacteria (Duarte et al., 2012), meningococci (Nieuwland et al., 2000), and staphylococci (Timar et al., 2013). However, only a few stimulator of NDTR generation were found. fMLP and TNF-α are the only known stimulators of NDTR formation *in vitro* (Lim et al., 2015), and NDTRs are found in *in vivo* models of influenza virus infection (Lim et al., 2015) and bacterial infections (Hyun et al., 2012). However, contrary to our expectation, most immunological stimuli could induce both NDTR and NDMV formation. These results suggest that the generation of NDTRs and NDMVs is not determined by stimulators that neutrophils encounter during the inflammatory process. Therefore, we next investigated the differences in generation mechanism between NDTRs and NDMVs. Although neutrophil-derived EVs shared common mechanisms of generation (e.g., via ERK MAPK, p38 MAPK, Rho, Rac1, Cdc42, and extracellular Ca^2+^), we found that integrin signaling and the PI3K pathway were required for the generation of NDTRs and NDMVs, respectively (Figure 1D). Since NDTRs are generated from neutrophils during diapedesis from blood vessels toward inflammatory foci (Hyun et al., 2012; Lim et al., 2015), the adhesion molecules highly expressed in endothelial cells ensure the generation of NDTRs by neutrophils. In contrast, neutrophils generate NDMVs in an environment devoid of adhesion molecules. Therefore, these results suggest that the generation of NDTRs and NDMVs by neutrophils is dependent on the immunological environment, not the type of stimulation.

We next tried to identify the specific markers of NDTRs and NDMVs. Previous studies reported various markers of NDMVs: phosphatidylserine (Dalli et al., 2008; Eken et al., 2013; 2010; Gasser, 2004; Kambas et al., 2014; Pitanga et al., 2014; Pliyev et al., 2014; Slater et al., 2017; Stein and Luzio, 1991; Timar et al., 2013), Annexin A_1_ (Dalli et al., 2008; Headland et al., 2015), neutrophil granules (Gasser et al., 2003; Gasser and Schifferli, 2005; Headland et al., 2015; Hess et al., 1999; Y. Hong et al., 2012; Mesri and Altieri, 1999; Nieuwland et al., 2000; Pitanga et al., 2014; Prakash et al., 2012), adhesion molecules (Dalli et al., 2008; Gasser et al., 2003; Y. Hong et al., 2012; Pitanga et al., 2014; Pluskota et al., 2008; Prakash et al., 2012; Slater et al., 2017), and complement receptors (Gasser et al., 2003; Gasser and Schifferli, 2005; Hess et al., 1999). In contrast, very little is known about specific markers of NDTRs, except for integrin-associated molecules such as lymphocyte function-associated antigen 1 (LFA-1) and MAC-1 (Hyun et al., 2012; Lim et al., 2015). Neutrophil-derived EVs shared most of markers. Both NDTRs and NDMVs expressed general EV markers (e.g., AnnV, AnnA1, CD9, and CD81), neutrophil-specific markers (e.g., CD63, CD66b, and CD35), and adhesion molecules (e.g., CD11b, CD49d, CD29, and CD18); in addition, both contained certain functional molecules (e.g., HSP and MCP-1). However, the expression level of CD16 distinguished NDTRs from NDMVs. NDMVs expressed relatively high levels of CD16 compared to those on NDTRs. CD16, an Fcγ type III receptor, is found on the surface of neutrophils (van de Winkel and Capel, 1993) and is preferentially distributed on the leading edge in migrating neutrophils (Seveau et al., 2001). NDTRs are generated from elongated uropods of migrating neutrophils; hence, the contribution of this physical characteristic to the generation mechanism might be responsible for the relatively low expression of CD16 on NDTRs.

Notably, the expression levels of some markers of neutrophil-derived EVs varied according to the stimulators. Although stimulation with TNF-α, C5a, and PMA induced marker expression patterns similar to those produced by fMLP, bacterial stimulation induced a marked reduction in the expression of almost all kinds of surface markers. Moreover, TGF-β stimulation induced different patterns of surface marker expression in neutrophil-derived EVs. For example, CD11b expression was nearly abrogated in TGF-β-induced neutrophil-derived EVs, and CD16 expression levels were reciprocally increased in NDTRs. These changes according to specific stimuli suggest that neutrophils might generate different types of EVs according to the immune environment. In support of this hypothesis, previous studies identified the functional changes in NDMVs in response to the stimulation. Bacteria-induced NDMVs gained the ability to inhibit bacterial growth, whereas spontaneously generated NDMVs had no bactericidal activity (Timar et al., 2013). Moreover, NDMVs generated from mycobacteria-infected neutrophils inhibited macrophage activation, while spontaneously generated NDMVs did not significantly affect macrophage function (Duarte et al., 2012). Therefore, studying the function and phenotype of different neutrophil-derived EVs according to the different types of stimulation could be interesting.

To further investigate the functional differences between neutrophil-derived EVs, the bactericidal activity of these EVs was assessed. Both NDTRs and NDMVs directly killed bacteria via ROS- and granule-dependent mechanisms (Figure 2). Although a previous study showed that NDMVs limit bacterial growth by inducing bacterial aggregation {Timar: 2013da}, the specific mechanism underlying the bactericidal activity of NDMVs is incompletely understood. Previous studied have reported that NDMVs can actively generate ROS {Pitanga: 2014kp} {Hong: 2012de} and contain neutrophil granules such as myeloperoxidase (Gasser et al., 2003; Y. Hong et al., 2012; Slater et al., 2017), lactoferrin, elastase, and proteinase (Gasser et al., 2003; Hess et al., 1999; Y. Hong et al., 2012; Mesri and Altieri, 1999). Therefore, ROS generation and protease-enriched granules are considered to be possible mechanisms of the direct bactericidal activity of NDMVs. Consistent with this hypothesis, NDMVs actively generate ROS in response to stimulation (Supplementary Figure 2E), and NADPH oxidase inhibitor significantly attenuated the bactericidal activity of NDMVs (Figure 2D and 2E). Moreover, NDMVs expressed granule markers (Figure 1E), and protease inhibitors significantly attenuated the bactericidal activity of NDMVs (Figure 2D and 2E). Interestingly, NDTRs also showed bactericidal activity via ROS- and granule-dependent mechanisms (Figure 2C). Additionally, NDTRs actively generated ROS in response to PMA stimulation (Supplementary Figure 2E) and expressed various granule markers (Figure 1E). Moreover, the bactericidal activity of NDTRs was significantly attenuated by the NADPH oxidase inhibitor and protease inhibitors (Figure 2D and 2E), suggesting that NDTRs not only guide the migration of immune cells to inflammatory foci but also provide a minimal defense against pathogens. Moreover, our results suggest that neutrophil-derived EVs might be involved in the general bactericidal activity of neutrophils (Figure 2C). After generating EVs, neutrophils exhibited a marked decrease in overall bactericidal activity, suggesting the development of an exhaustion state (Figure 2C). Neutrophils are exposed to various bacteria-derived products which are strong stimulators of neutrophil-derived EVs production. Therefore, neutrophils might be forced to produce EVs during classical phagocytosis-mediated bactericidal processes, and the bactericidal activity of neutrophil-derived EVs might be involved in the general bactericidal activity of neutrophils.

The primary function of EVs is to mediate intercellular communications (Camussi et al., 2010; Théry et al., 2009; van der Pol et al., 2012), and neutrophil-derived EVs also mediate cell-to-cell communications between neutrophils and neighboring cells. The results of previous studies indicated that NDMVs induce anti-inflammatory functions in neighboring cells. NDMV-exposed NK cells exhibited increased expression of anti-inflammatory cytokines (Pliyev et al., 2014), and NDMVs inhibited the maturation of monocytes into dendritic cells (Pliyev et al., 2014). Moreover, NDMVs inhibit proinflammatory cytokine expression in monocytes, macrophages (Eken et al., 2013; 2010; Prakash et al., 2012; Rhys et al., 2018), and endothelial cells (Mesri and Altieri, 1999; 1998). In contrast, very little is known regarding the effects of NDTRs in cell-to-cell communications except that they guide following CD8^+^ T cells and monocytes toward inflammatory foci (Lim et al., 2015). We found that neutrophil-derived EVs play a pivotal role in recruiting monocytes and skewing the differentiation of monocytes into either proinflammatory or anti-inflammatory macrophage. Both subtypes of neutrophil-derived EVs not only induced monocyte chemotaxis (Figure 3) and but also phagocytosed by monocytes (Figure 4A and 4B). Phagocytosed neutrophil-derived EVs showed differential effects on the phenotypic polarization of macrophages; NDTRs induced polarization toward a proinflammatory phenotype, whereas NDMVs induced polarization toward an anti-inflammatory phenotype (Figure 4C-E).

We found that miRNAs differentially expressed in NDTRs and NDMVs are responsible for macrophage polarizations into different phenotypes. Interestingly, most miRNAs found in NDTRs are associated with proinflammatory responses of macrophages, whereas most miRNAs found in NDMVs are associated with anti-inflammatory responses of macrophages. miR-1260a and miR-1285-5p are expressed in bacteria-infected macrophages (Derrick et al., 2017; 2013; Meng et al., 2014). miR-122-5p and miR-29a-3p induce proinflammatory gene expression in macrophages (Fabbri et al., 2012; Momen-Heravi et al., 2015). Moreover, miR-24-3p and miR-4454 are found in monocyte-derived dendritic cells (Fell et al., 2016) and alveolar macrophages (Armstrong et al., 2017). In contrast, miR-126-3p and miR-150-5p are expressed in M2 polarized macrophages (Cobos Jiménez et al., 2014; Escate et al., 2016) and are associated with suppression of inflammation (Harris et al., 2008; Lin et al., 2018; Manoharan et al., 2014; Shantikumar et al., 2012). Our results showed differential miRNA expression in NDTRs and NDMVs; thus, neutrophils incorporate different types of miRNAs into EVs according to the immune environments. Since the adhesion molecules differentiate the generation mechanism of NDTRs and NDMVs, signaling associated with adhesion molecules could be an explanation for this phenomenon.

The reason that neutrophils generate different types of EVs is unknown. However, our findings suggest that neutrophils produce different types of EVs according to the immune environments to orchestrate inflammation. Successful inflammation requires effective initiation and resolution. Since neutrophils are the first cells recruited to sites of inflammation, they play a pivotal role in initiating inflammation (Amulic et al., 2012; Nauseef, 2016; Nauseef and Borregaard, 2014). Neutrophils actively participate in eliminating pathogens, guide following immune cells by releasing various kinds of chemokines and cytokines, and modulate the functions of neighboring immune cells (Nauseef, 2016; Soehnlein et al., 2017). Neutrophils also play an important role in the resolution of inflammation; they can persist at inflammatory foci during the entire process of inflammation and participate in the resolution of inflammation by releasing proresolving factors (Serhan et al., 2008). As discussed above, the most prominent difference between NDTRs and NDMVs is the spatiotemporal generation mechanism; NDTRs are generated from neutrophils migrating toward inflammatory foci, whereas NDMVs are generated from neutrophils that have arrived at inflammatory foci. Therefore, NDTRs might augment the proinflammatory responses of accompanying immune cells to combat imminent inflammatory insults, whereas NDMVs might limit excessive inflammation by enhancing the anti-inflammatory responses of immune cells at inflammatory foci. In support of this idea, NDTRs demonstrated a profound beneficial effect in murine models of inflammation. NDTRs reduced lethality in the murine model of sepsis (Figure 6A) and pathological changes in the murine model of chronic colitis (Figure 6B-D). Moreover, NDTRs increased the number of proinflammatory macrophages in both murine inflammatory models. Therefore, we believe that neutrophils generate NDTRs during migration toward inflammatory foci to orchestrate the initiation of inflammation. In contrast, NDMVs did not demonstrate any beneficial effects in these murine models of inflammation, suggesting negligible effects of NDMVs on the overall outcome in our *in vivo* inflammation models. However, previous study showed the beneficial effect of NDMVs in chronic inflammation. The intra-articular injection of NDMVs decreased cartilage damage in a murine model of inflammatory arthritis by enhancing TGF-β production from chondrocytes (Headland et al., 2015). Since our study showed that NDMVs promote M2 macrophage polarization, it could be additional mechanism underlying the beneficial effect of NDMVs on inflammatory arthritis. However, additional *in vivo* studies are needed to verify the anti-inflammatory effects of NDMVs.

In conclusion, our study provides important insight into the differential functions of neutrophil-derived EVs; proinflammatory NDTRs and anti-inflammatory NDMVs. NDTRs guide monocytes and induce polarization toward proinflammatory macrophages, thereby contributing to the effective initiation of inflammation. On the other hand, NDMVs have similar characteristics to NDTRs, but they induce macrophage polarization into an anti-inflammatory phenotype. Therefore, modulating neutrophil-derived EV generation might be a new strategy for modulating inflammation.

## EXPERIMENTAL PROCEDURES

### Neutrophil Isolation

Human blood experiment was approved by Institutional Research Board of Kyungpook National University and Hallym University. Neutrophils were isolated as previously described (C.-W. Hong et al., 2010).

### Live cell imaging of NDMV and NDTR formation

Neutrophils were stained with cell tracker and seeded in either confocal dish (SPL Life Sciences) for evaluation of NDMVs or μ-slide chamber (ibidi) pre-coated with fibronectin (5 μg/ml, Merck Millipore) for evaluation of NDTRs. Cells were treated with various stimulators for neutrophil EVs generation, and were visualized by immunofluorescence microscopy (Olympus IX83, Olympus). Neutrophils were stimulated with various stimulants: fMLP (1 μM, Sigma-Aldrich), LPS (1 μg/ml, Sigma-Aldrich), C5a (50 ng/ml, Sino Biologicals), S100B (100 ng/ml, Sino Biologicals), HMGB1 (100 ng/ml, Sino Biologicals), TNF-α (50 ng/ml, Sino Biologicals), IFN-γ (100 ng/ml, Sino Biologicals), TGF-β (20 ng/ml, Sino Biologicals), IL-4 (20 ng/ml, Sino Biologicals), PMA (100 μg/ml, Sigma-Aldrich), and L-NAME (10 μM; Tocris Bioscience).

### Isolation of NDMV and NDTR

For isolation of NDMVs, supernatants were collected from stimulated neutrophils on uncoated culture plates. For isolation of NDTRs, neutrophils were stimulated on culture plate coated with fibronectin (5 μg/ml) and adherent NDTRs were recovered by cell scraper. The purification of neutrophil EVs were performed by centrifugation at 2500 rpm and filtration through 1.2 μm filter (Ministart Syringe Filter, Sartorius). The filtered EV-containing supernatants were ultra-centrifuged at 100,000 × g for 60 min at 4 •. The pelleted NDTRs and NDMVs were dissolved in 100 μl phenol red-free RPMI and stored at -70 •.

### Quantification of NDMV and NDTR

EVs were isolated from neutrophils pre-stained with calcein-AM (20 μg/ml, Merck Millipore). The calcein-AM fluorescence in neutrophil EVs were measured using Spectramax M2/e fluorescence microplate reader (Molecular Devices). For inhibition of neutrophil EVs generation, neutrophils were stimulated with fMLP (1 μM) in presence of various inhibitors: calcium chelator (BAPTA-AM, 10 μM, Tocris), inhibitor of calcium-activated chloride channel (CaCCinh-A01, 30 ng/ml, Sigma-Aldrich), inhibitor of Rho-A GTPase (Tat C3, 20 ng/ml, provided from Prof. J. Park in Hallym University), Inhibitor of Rac-1 GTPase (NSC23766, 10 μM, Tocris bioscicence), inhibitor of actin polymerization (Cytochalasin D, 50 μg/ml, Sigma-Aldrich), inhibitor of PI3-Kinase (Wortmanin, 100 nM, Tocris), inhibitor of p38 MAP kinase (SB203580, 10 μM, Tocris), inhibitor of ERK MAP kinase (PD90859, 10 μM, Tocris), inhibitor of Cdc42 (ML141, 10 μM, Tocris), inhibitor of β1 integrin very late antigen (VLA-4) (Bio1211, 10 μM, Tocris bioscicence), inhibitor of macrophage-1 (MAC-1) antigen, IMB10 (10 μM, Sigma-Aldrich), inhibitors on the autophagic flux pathways (3-MA, 5 μM, Sigma-Aldrich; chloroquine, 5 μM, Sigma-Aldrich; bafilomycin, 5 μM, Sigma-Aldrich), and inhibitors on the mTOR pathway (Rapamycin, 10 nM, Sigma-Aldrich).

### Phenotyping of NDMV and NDTR using flow cytometry

Isolated EVs were fixed and stained with Annexin V (FITC, Abcam), FITC-conjugated Annexin A1 (Annexin A1, Bio Legend; FITC conjugate, Abcam), CD9 (FITC, Bio Legend), CD81 (FITC, Bio Legend), CD63 (FITC, BD biosciences), CD66b (FITC, BD biosciences), CD35 (PE, BD bioscience), CD11b (FITC, Invitrogen), CD49d (FITC, Bio Legend), CD29 (PE, eBioscience), CD18 (PE, eBioscience), and CD16 (PE, BD biosciences). For intracellular staining, fixed neutrophil-derived EVs were permeabilized with phosflow perm buffer (BD biosciences), and stained with MCP (FITC, eBiosciences) and HSP (APC, Invitrogen).

### Nanoparticle tracking analysis (NTA)

Isolated EVs were re-suspended in PBS and further diluted for NTA analysis with the Nanosight LM10 nanoparticle characterization system (Malvern Instruments). NTA analytical software version 3 was used for capturing and analyzing data.

### Bactericidal activity

Opsonized *E. coli* (DH5α) and *S. aureus* (ATCC 25923) were exposed to EVs isolated from neutrophils stimulated with respective bacteria. For inhibition of bactericidal pathways, bacteria were exposed to EVs in presence of either protease inhibitor cocktail (Sigma-Aldrich), diphenylene iodonium (DPI, an inhibitor of reduced nicotinamide adenine dinucleotide phosphate oxidase, 10 μM, Molecular Probes), or DNase I (10 μM, Bio basic Canada Inc.). For bactericidal activities of remnant neutrophils, neutrophils were allowed to generate neutrophil EVs in response to respective bacteria and the bactericidal activities of remnant neutrophils were measured.

### Monocyte Isolation

Monocytes were obtained from peripheral blood mononuclear cells (PBMCs) layer obtained after histopaque centrifugation by using percoll solution as described previously (Repnik et al., 2003). Macrophages were differentiated from monocytes by incubation with granulocytes-macrophage colony stimulating factor (GM-CSF, 10 ng/ml, Bio Legend)

### Chemotaxis assay

Migration of monocytes and macrophages against NDMV and NDTR were determined using μ-slide chamber (ibidi) following manufacturer’s instruction. The cell movement were visualized and captured by Nikon Eclipse Ni-U microscope using 20X objective. Cell movement was then analyzed using ImageJ (Schneider et al., 2012) and Chemotaxis and migration tool (ibidi).

### Polarization of differentiated THP-1

THP-1 cells (Korean Cell Line Bank) were differentiated into M0 macrophages by treatment with PMA (100 ng/ml). M0-differentiated THP-1 cells were further exposed to neutrophil EVs which were isolated from fMLP-stimulated neutrophils. For control, M1 macrophages were induced by LPS (100 ng/ml) and IFN-γ (10 ng/ml, Sinobiologicals), and M2 macrophages were induced by IL-4 (10 ng/ml, Sinobiologicals) and IL-12 (10 ng/ml, Sinobiologicals). Phenotypic markers for FACS analysis include CD14 (PerCP, BD biosciences), HLA-DR (APC, BD biosciences), CD80 (PE, Bio Legend), CD86 (PE, Bio Legend), CD209 (APC, Bio Legend), CD23 (APC, Bio Legend), CD163 (PE, Bio Legend), CD206 (FITC, BD biosciences), and CD1a (FITC, Bio Legend). For evaluation of cytokine expressions, RNA was extracted from THP-1 and cDNA was synthesized using RT^2^ First strand kit (Qiagen). Pre-designed RT^2^ qPCR primer assays (Qiagen) were used to determine the expression level of cytokines such as: iNOS (NOS2, PPH00173F), Arg-1 (PPH20977A), TNF-α (PPH00341F), TGF-β1 (PPH00508A), IL-10 (PPH00572C), IL-12A (PPH00544B), IL-6 (PPH00560C) and IL-1β (PPH00171C).

### THP-1 uptake NDMV and NDTR

Isolated NDTRs and NDMVs from neutrophils pre-stained with cell tracker green (5 μg/ml) were exposed to M0-differentiated THP-1 cells on confocal dish. The cell movements were visualized using immunofluorescence microscopy at 37 •, 5% CO_2_ for 1 h.

### miRNA array and validation

RNA was isolated from EVs using miRNeasy Micro Kit (Qiagen) and cDNA was synthesized with TaqMan MicroRNA reverse transcription kit (Applied Biosystems). Reactions were performed using a TaqMan Universal Master Mix II (Applied Biosystems) and TaqMan probe (hsa-miR-24-3p, hsa-miR-29a-3p, hsa-miR-122-5p, hsa-miR-126-3p, hsa-miR-150-5p, hsa-miR-451a, hsa-miR-1260a, hsa-miR-1285-5p, hsa-miR-4454, hsa-miR-7975). Cel-miR-39 was added as an exogenous control for normalization. The miRNA profile of samples was analyzed with Nanostring nCounter™ system. To validate the effects of miRNAs on the phenotype polarization of macrophages, M0-differentiated THP-1 cells were transfected with miRNA mimics (hsa-miR-1285-5p, hsa-miR-1505p, hsa-miR-451a, hsa-miR-1260a, hsa-miR-4454, hsa-miR-7975, and hsa-miR-126-3p, abm) using NEPA21 (NEPA gene). RNA was extracted and the expression of iNOS and arginase was measured using quantitative PCR.

### Animal experiments

Animal experiments were approved by the Institutional Animal Care and Use Committee of Kyungpook National University and Hallym University. BALB/c (male, 5-8 weeks old) mice were purchased from Orient Bio. Mouse neutrophils were isolated from bone marrow of healthy mouse using neutrophil enrichment kit (Miltenyi Biotec) or EasySep™ mouse neutrophil enrichment kit (StemCell technologies) as previously described (Park et al., 2017). EVs were isolated from neutrophils stimulated with *E. coli*. Experimental sepsis was induced in BALB/c mice by the cecal ligation and puncture (CLP) procedure as previously described (Yan et al., 2004). BALB/c mice were treated with either NDTRs (100 μl, i.p.), NDMVs (100 μl, i.p.) or vehicle (saline, 100 μl, i.p.) 30 min before CLP surgery, and further treated on day 1, 2, and 3 after CLP surgery. The survival of septic mice were monitored for 9 days. DSS-induced chronic colitis was performed according to previous study (Marcon et al., 2013). In brief, BALB/c mice received water containing 2% DSS during first 5 days and further received normal drinking water for 10 days. At day 16, mice were exposed to water containing 2% DSS again until day 22. Mice were treated with NDTRs and NDMVs intraperitoneally on days 16, 18, 20, and 22.

### Histological analysis

Colon were removed at day 22 after DSS-induced colitis, fixed with 4% formaldehyde solution (Biosesang), embedded in paraffin, and stained with hematoxylin (Dako Mayer’s Hematoxylin, Agilent) and eosin. (Daejung chemical and metal Co Ltd.). Histological analysis were performed as previously described (Marcon et al., 2013). In brief, epithelial hyperplasia, mononuclear cells and polymorphonuclear cell infiltration in lamina propria, crypt inflammation, epithelial hyperplasia, and erythrocyte loss were scored‥

### Serum evaluation

Serum obtained from sepsis patients who admitted to the intensive care unit of Kyungpook National University Hospital between April and October 2016. All patients provided informed consents in accordance with the Declaration of Helsinki. Serum were obtained from healthy volunteers or sepsis patients and further were filtered through 1.2 µM filter. The filtrate were then treated with 5µl of ExoQuick (System Biosciences), incubated for 30 mins at 4 □, followed by high-speed centrifugation (11,600 rpm for 1 hr). The pellets were then fixed and stained with Annexin V (FITC) and CD16 (PE) antibody.

### Statistical analysis

Data are presented as the mean ± SEM for continuous variables and as the number (%) for the categorical variables. Comparisons between two groups were performed with either two-tailed Student’s t test (parametric) or Mann-Whitney (non-parametric test). Survival data were analyzed using Mantel-Cox log-rank test. Values of P < 0.05 were considered to indicate statistical significance. Statistical data were analyzed by Graphpad prism 7.0 (GraphPad Software Inc.).

A detailed description of all methods is available in **supplementary information**.

## Abbreviations

EV: extracellular vesicle
NDMV: neutrophil-derived microvesicles
NDTR: neutrophil-derived trails
NK cells: natural killer cells
MoDCs: monocyte-derived dendritic cells
CXCL12: CXC-chemokine ligand 12
PAMPs: pathogen-associated molecular patterns
fMLP: formyl-methionyl-leucyl-phenylalanine
LPS: lipopolysaccharide
*E. coli*: *Escherichia coli*
*S. aureus*: *Staphylococcus aureus*
DAMPs: danger-associated molecular patterns
HMGB1: high mobility group box 1
TNF-α: tumor necrosis factor-α
IFN-γ: interferon-γ
TGF-β: tumor growth factor-β
IL-4: interleukin-4
L-NAME: N(γ)-nitro-L-arginine methyl ester
PMA: phorbol 12-myristate 13-acetate
MCP-1: monocyte chemoattractant protein 1
iNOS: inducible nitric oxide synthase

