## Supplementary file for "Neutrophil-derived extracellular vesicles: proinflammatory trails and anti-inflammatory microvesicles"

**SUPPLEMENTAL INFORMATION**

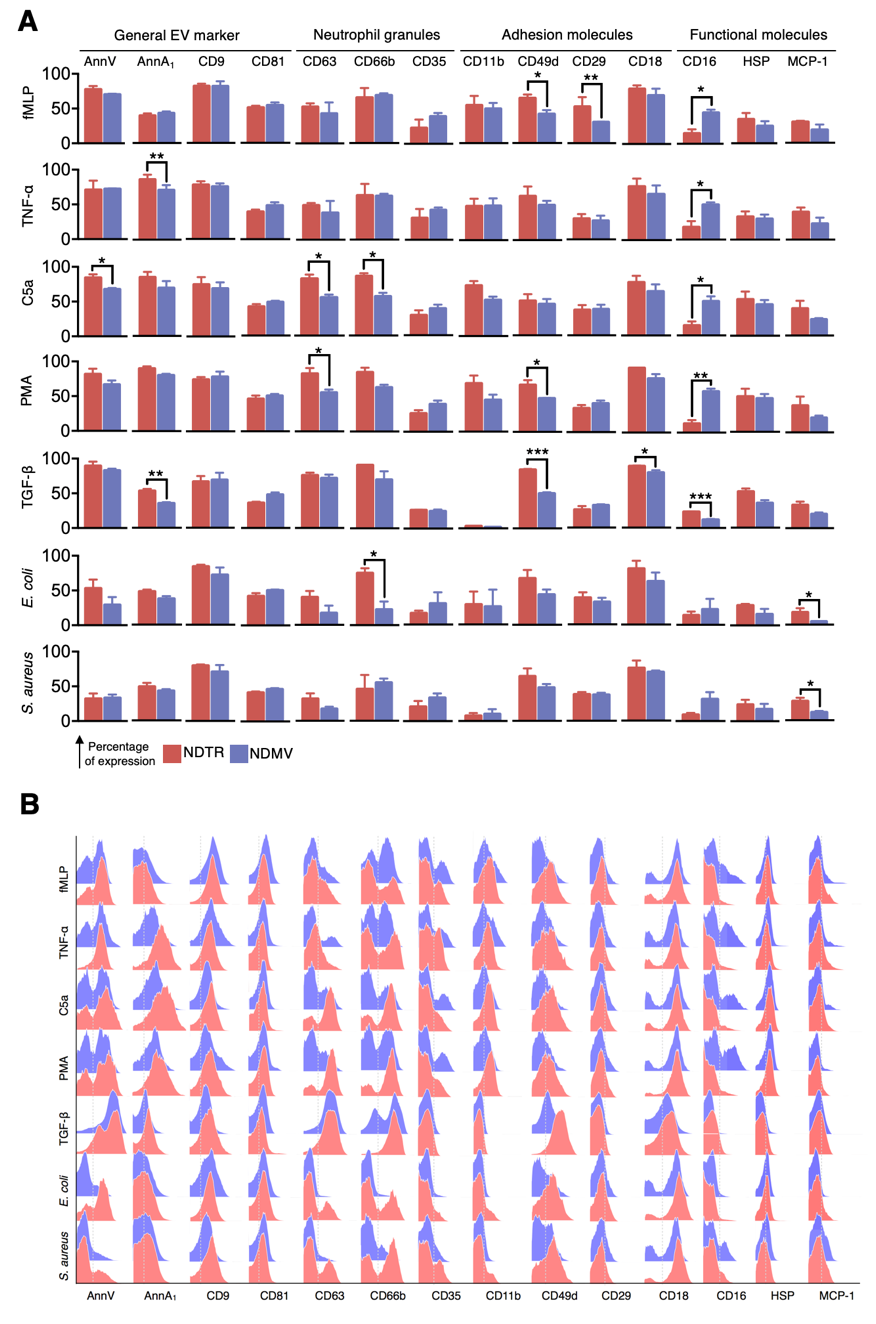

**Figure S1. Related to Figure 1. Phenotype characterization of neutrophil-derived EVs.** NDTRs and NDMVs were isolated from neutrophils stimulated with indicated stimulators. Neutrophil-derived EVs were gated based on the forward- and side-scatter profiles. The surface expression of Annexin V, Annexin A1, CD9, CD81, CD63, CD66b, CD35, CD11b, CD49d, CD29, CD18, and CD16 were examined. For evaluation of HSP and MCP-1, neutrophil-derived EVs were permeabilized and expression levels of indicated markers were examined. (A) The percentage of expressions. (B) Representative histogram. All data are representative of more than three independent experiments.

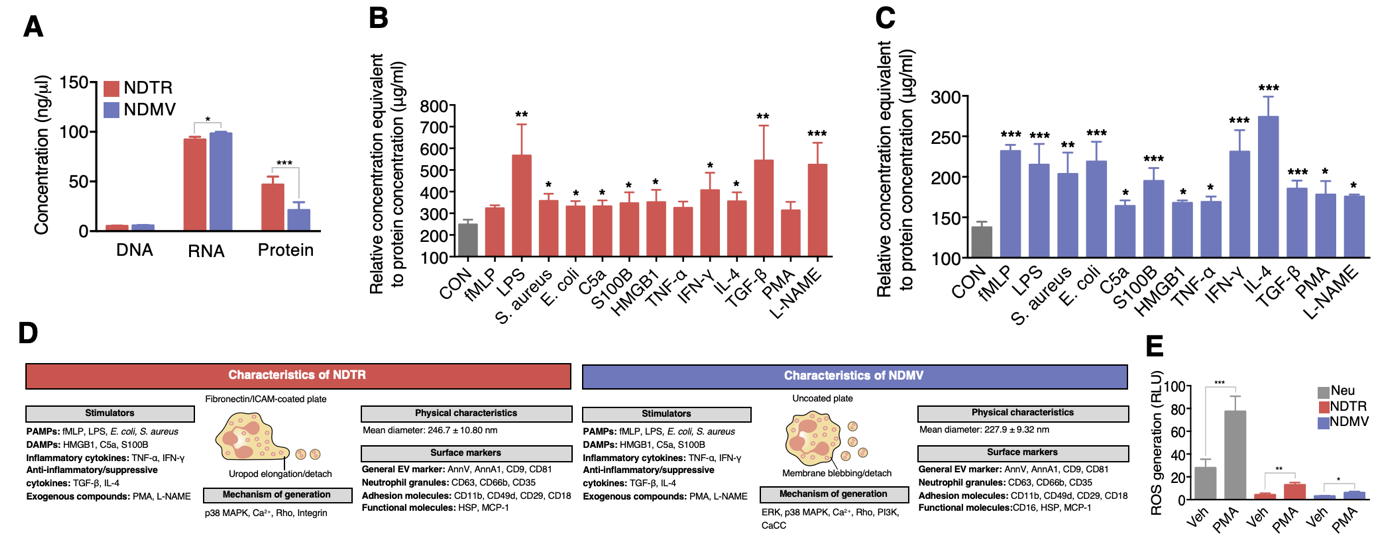

**Figure S2. Related to Figure 1 (A-D) and 2 (E). Validation of contents in neutrophil-derived EVs.** (A) Quantification of contents in neutrophil-derived EVs. EVs were isolated from fMLP-stimulated neutrophils. n = 3. (B-C) Quantification of protein concentration of neutrophil-derived EVs according to indicated stimulators. Neutrophil EVs were isolated from Calcein-AM-stained neutrophils with the indicated stimulators, and fluorescence in neutrophil-derived EVs were calculated to equivalent protein concentration. n = 3. (D) Schematic illustration of neutrophil EV characterization. (E) ROS generation from neutrophil-derived EVs. Isolated neutrophil-derived EVs were stained with DCF-DA and stimulated with PMA for 30 min. n = 3. The data are shown as the mean ± SEM. *P < 0.05; **P < 0.01; ***P < 0.001.

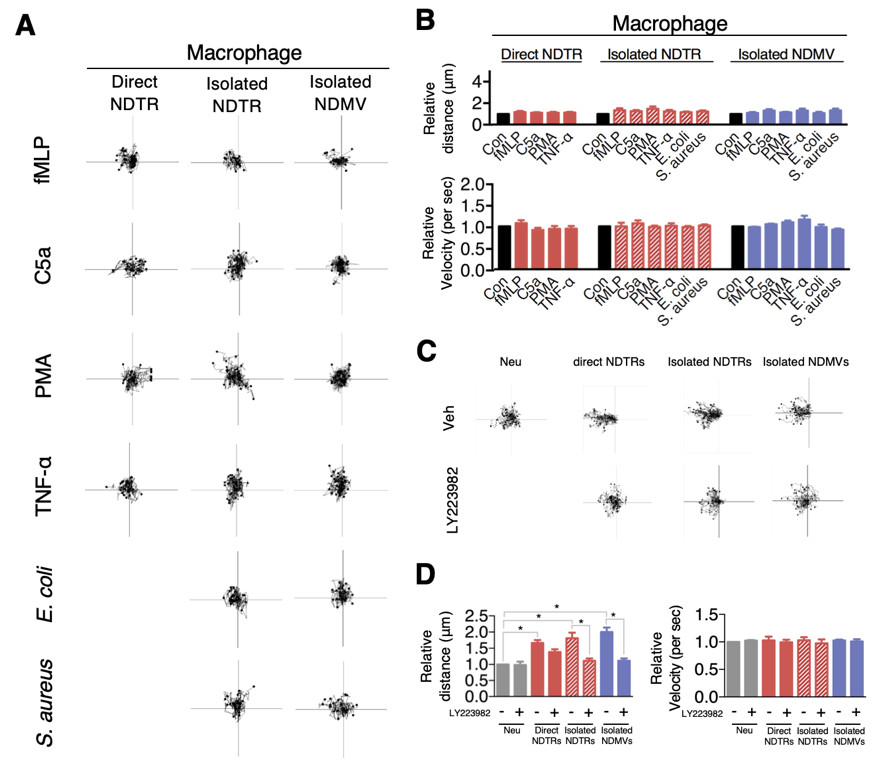

**Figure S3. Related to Figure 3. Chemotaxis of macrophages and neutrophils against neutrophil-derived EVs.** (A) Tracking analysis on macrophage chemotaxis against neutrophil-derived EVs. The distances of migrated cells were tracked on every minute for 45 min. The representative tracking results of thirty cells per each group were presented. (B) Relative mean distance and the relative mean velocity of macrophages toward neutrophil-derived EVs. (C) Tracking analysis on the neutrophil chemotaxis. (D) Relative mean distance and velocity of neutrophils toward neutrophil-derived EVs. LY223982: neutrophils were loaded in the presence of BLT_1_ receptor antagonist. Direct NDTR: neutrophils were allowed to generate NDTRs according to indicated stimulators on a fibronectin-coated μ-slide chamber and target cells were allowed to move directly toward NDTRs. Isolated NDTRs and NDMVs: neutrophils were stimulated with indicated stimulators and EVs were isolated. The lane of μ-slide chamber was coated with isolated neutrophil-derived EVs and target cells were allowed to move toward neutrophil EVs. The data are shown as the mean ± SEM. *P < 0.05; **P < 0.01; ***P < 0.001. All data are representative of three independent experiments (n = 3).

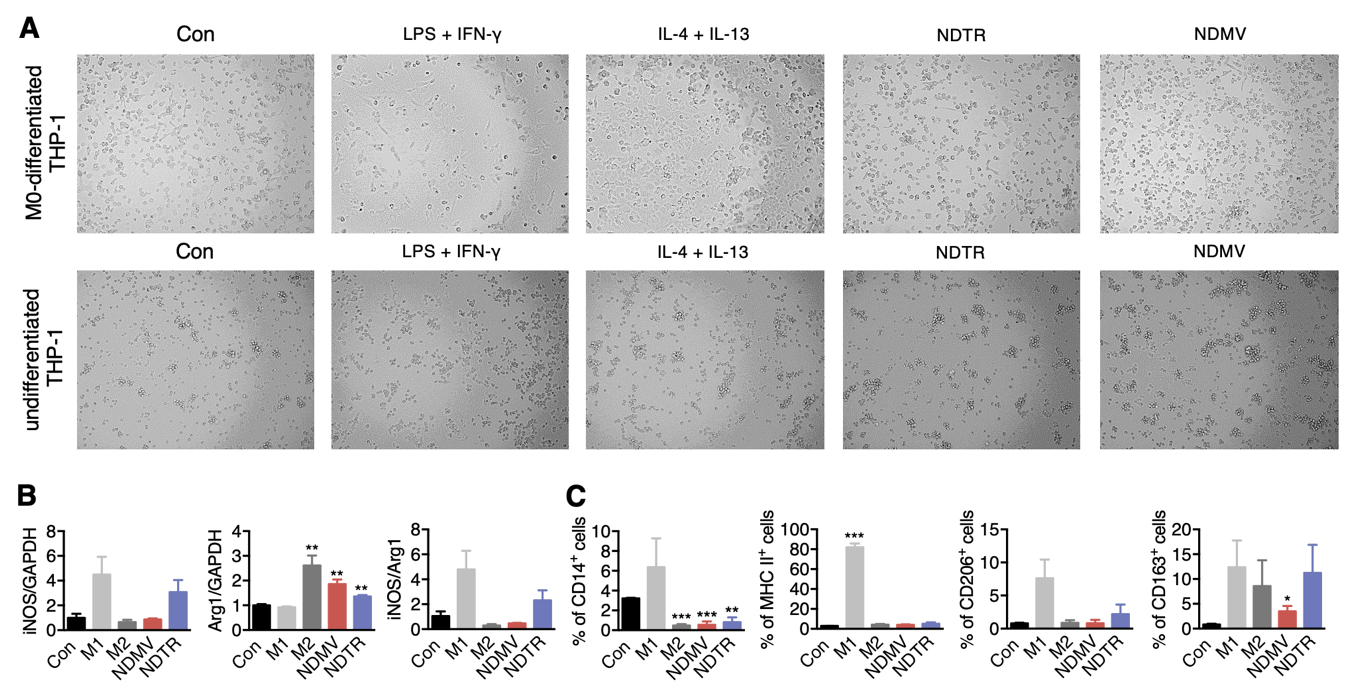

**Figure S4. Related Figure 4. Effects of neutrophil-derived EVs on phenotype polarization of undifferentiated THP-1 cells.** (A) Morphology of M0-differentiated and undifferentiated THP-1 cells. (B-C) The expression of M1 and M2 macrophage-associated markers in undifferentiated cells exposed to neutrophil-derived EVs (n = 3 per each group.) The data are shown as the mean ± SEM. *P < 0.05; **P < 0.01; ***P < 0.001 compared to control.

**Movie S1, related Figure 1.** Live imaging of NDTR generation from neutrophils. Neutrophils stained with call tracker were incubated with μ-slide chamber pre-coated with fibronectin in presence with various stimulators.

**Movie S2, related Figure 1.** Live imaging of NDMV generation from neutrophils. Neutrophils stained with call tracker were incubated with confocal plates in presence with various stimulators.

**Movie S3, related Figure 4.** Live imaging of NDTR uptake by M0-differentiated THP-1 cells. NDTRs were stained with calcein-AM

**Movie S4, related Figure 4.** Live imaging of NDMV uptake by M0-differentiated THP-1 cells. NDMVs were stained with calcein-AM

**Material and methods**

**Neutrophil Isolation**

Human blood experiment was approved by Institutional Research Board of Kyungpook National University and Hallym University. Venous blood was taken from healthy volunteers. Neutrophils were purified using histopaque (Sigma-Aldrich) centrifugation followed by Dextran sedimentation as described previously {Hong:2010ca}. Briefly, human venous blood was drawn from healthy male volunteers into vials containing anti-coagulant, layered over an equal volume of histopaque 1077 (Sigma-Aldrich), and centrifuged at 2500 rpm for 30 mins at RT. The neutrophil-containing lower layer was collected and sedimented with 5% (w/v) dextran (Pharmacosmos, Holbaek, Denmark) for 45 min at 4 ℃. Neutrophil-rich layer was collected and remaining red blood cells were removed using hypotonic lysis. The cells were finally resuspended in RPMI 1640 (Gibco) supplemented with 5% Fetal bovine serum (FBS, HyClone) at 1 × 10^7^/ml. The purity of neutrophils was determined by Wright-Geimsa staining. The purity was consistently greater than 95%. All the experiments regarding NDTRs and NDMVs were conducted in RPMI unless otherwise indicated.

**Live cell imaging of NDMV and NDTR formation**

For evaluation of NDMV formation, neutrophils (5 × 10^6^ cells) were stained with cell tracker green (1.5 - 2.0 μg/ml, Invitrogen) for 30 min and 3 × 10^6^ cells were seeded in confocal plates (SPL Life Sciences). For evaluation of NDTR formation, neutrophils (5×10^6^ cells) were stained cell tracker green for 30 min and 3 × 10^6^ cells were incubated in the μ-slide chamber (Ibidi) pre-coated with fibronectin (5 μg/ml, Merck Millipore). Then neutrophils were stimulated with various stimulators. The released NDMV and NDTR were visualized by immunofluorescence microscopy (Olympus IX83, Olympus). Visualization of the cells were carried out at 37 ℃, 5% CO_2_ for 1 hour. Neutrophils were stimulated with various stimulators: fMLP (1 μM, Sigma-Aldrich), LPS (1 μg/ml, Sigma-Aldrich), C5a (50 ng/ml, Sino Biologicals), S100B (100 ng/ml, Sino Biologicals), HMGB1 (100 ng/ml, Sino Biologicals), TNF-α (50 ng/ml, Sino Biologicals), IFN-γ (100 ng/ml, Sino Biologicals), TGF-β (20 ng/ml, Sino Biologicals), IL-4 (20 ng/ml, Sino Biologicals), PMA (100 μg/ml, Sigma-Aldrich), and L-NAME (10 μM; Tocris bioscicence).

**Isolation of NDMV and NDTR**

For isolation of NDMVs, neutrophils (5 × 10^6^ cells) were seeded on the culture plate and stimulated with stimulators for 20 min. Supernatants were collected from uncoated plates and were subjected to purification. For isolation of NDTRs, neutrophils (5 × 10^6^ cells) were seeded on the culture plates coated with fibronectin (5 μg/ml, Merck Millipore) and stimulated with stimulators for 20 min. Neutrophils and media were discarded from fibronectin-coated plated and adherent NDTRs were recovered using cell scraper. The remnant neutrophils and debris were further removed by centrifugation at 2500 rpm and filtration through 1.2 μm filter (Minisart Syringe Filter, Sartorius). The filtered EV-containing supernatants were ultra-centrifuged at 100,000 × g for 60 min at 4 ℃. The pelleted NDTRs and NDMVs were dissolved in 100 μl phenol red-free RPMI and stored at -70 ℃.

**Quantification of NDMV and NDTR**

For quantification of NDTRs and NDMVs, neutrophil EVs were isolated from neutrophils stained with calcein-AM. Neutrophils (5 × 10^6^ cells) were stained with calcein-AM (20 μg/ml, Merck Millipore) and stimulated with various stimulators: fMLP (1 μM), LPS (1 μg/ml), *E. coli* (1 × 10^6^ ), *S. aureus* (1× 10^6^), C5a (50 ng/ml), S100B (100 ng/ml), HMGB1 (100 ng/ml), TNF-α (50 ng/ml), IFN-γ (100 ng/ml), TGF-β (20 ng/ml), IL-4 (20 ng/ml), PMA (100 μg/ml), and L-NAME (10 μM). Then neutrophil EVs were isolated and the fluorescence were measured using Spectramax M2/e fluorescence microplate reader (Molecular Devices). For inhibition of neutrophil EVs generation, neutrophils were stimulated with fMLP (1 μM) in presence of various inhibitors: calcium chelator (BAPTA-AM, 10 μM, Tocris bioscicence), inhibitor of calcium-activated chloride channel (CaCCinh-A01, 30 ng/ml, Sigma-Aldrich), inhibitor of Rho-A GTPase (Tat C3, 20 ng/ml, provided from Prof. Park in Hallym University), Inhibitor of Rac-1 GTPase (NSC23766, 10 μM, Tocris bioscicence), inhibitor of actin polymerization (Cytochalasin D, 50 μg/ml, Sigma-Aldrich), inhibitor of PI3-Kinase (Wortmanin, 100 nM, Tocris bioscicence), inhibitor of p38 MAP kinase (SB203580, 10 μM, Tocris bioscicence), inhibitor of ERK MAP kinase (PD90859, 10 μM, Tocris bioscicence), inhibitor of Cdc42 (ML141, 10 μM, Tocris bioscicence), inhibitor of β1 integrin very late antigen (VLA)-4 (Bio1211, 10 μM, Tocris bioscicence), inhibitor of macrophage-1 (MAC-1) antigen (IMB10, 10 μM, Sigma-Aldrich), inhibitors on the autophagic flux pathways (3-MA, 5 μM, Sigma-Aldrich; chloroquine, 5 μM, Sigma-Aldrich; bafilomycin, 5 μM, Sigma-Aldrich), and inhibitors on the mTOR pathway (Rapamycin, 10 nM, Sigma-Aldrich).

**Phenotyping of NDMV and NDTR using flow cytometry**

Isolated NDTRs and NDMVs were fixed with Phosflow buffer (BD biosciences) and stained with fluorescence tagged anti-human antibodies for 1 h at 4 ℃. For surface marker evaluation, neutrophil EVs were stained with Annexin V (FITC, Abcam), FITC-conjugated Annexin A1 (Annexin A1, Bio Legend; FITC conjugate, Abcam), CD9 (FITC, Bio Legend), CD81 (FITC, Bio Legend), CD63 (FITC, BD biosciences), CD66b (FITC, BD biosciences), CD35 (PE, BD biosciences), CD11b (FITC, Invitrogen), CD49d (FITC, Bio Legend), CD29 (PE, eBioscience), CD18 (PE, eBioscience), and CD16 (PE, BD biosciences). For evaluation of contents, fixed EVs were permeabilized with phosflow perm buffer (BD biosciences), and stained with MCP (FITC, eBioscience) and HSP (APC, Invitrogen). Neutrophil-derived EVs were identified using 1 μM beads (Life Technologies) and data acquisition was performed with BD FACS Calibur (BD Biosciences). The data was analyzed using FlowJo software (TreeStar Inc.).

**Nanoparticle tracking analysis (NTA)**

Isolated NDMV and NDTR were re-suspended in PBS and analyzed with the Nanosight LM10 nanoparticle characterization system (Malvern Instruments). Samples were manually injected into the sample chamber and each sample was measured in duplicate with an acquisition time of 30 sec and detection threshold setting 7. At least 975 frames were analyzed per video. NTA analytical software version 3 was used for capturing and analyzing data.

**Bactericidal activity**

Neutrophils (10^8^ cells) were stimulated with either opsonized E. coli (DH5α, 10^7^ cells, American Type Culture Collection, ATCC) or S. aureus (10^7^ cells, ATCC) for 30 min on the either uncoated culture plate or fibronectin-coated culture plate. Neutrophil-derived EVs were isolated according to the procedure as described earlier. Then opsonized E. coli (10^7^ cells) and S. aureus (10^7^ cells) were exposed to either NDTRs and NDMVs which were isolated from neutrophils stimulated with respective bacteria. After 30 min incubation, bacteria were washed and seeded on the agar plate. For comparison of bactericidal activity, bacteria were exposed to neutrophils (10^8^ cells) for 30 min, cells were lysed with chilled DDW, and remnant bacteria were seeded on agar plates. The percentages of killing were calculated as: percentages of killing (%) = [1 - (colony counts in experimental group/colony counts in vehicle (DDW)-treated group) × 100]. For inhibition of specific bactericidal pathways, bacteria were exposed to neutrophil EVs in presence of either protease inhibitor cocktail (Sigma-Aldrich), diphenylene iodonium (DPI, an inhibitor of reduced nicotinamide adenine dinucleotide phosphate oxidase, 10 μM, Molecular Probes) , or DNase I (10 μM, Bio basic). For bactericidal activities of remnant neutrophils, neutrophils (10^8^ cells) were allowed to generate either NDTRs or NDMVs in response to stimulation with respective bacteria. Then neutrophils were carefully recovered by removing of EVs using centrifugation at 2500 rpm and the bactericidal activities of recovered neutrophils (10^8^ cells) against bacteria (10^7^ cells) were examined. For generation of remnant neutrophils after generation of both NDTRs and NDMVs, neutrophils were first allowed to generate NDMVs in response to respective bacteria and neutrophils were recovered. Then neutrophils were allowed to generate NDTRs in response to respective bacteria and remnant neutrophils were recovered. Finally, bactericidal activity of remnant neutrophils were assessed.

**Monocyte Isolation**

Monocytes were obtained from peripheral blood mononuclear cells (PBMCs) layer obtained after histopaque centrifugation by using percoll solution as described previously {Repnik:2003ev}. Briefly, the PBMCs were layered on to hyper-osmotic percoll solution (GE healthcare Life Science). The obtained monocytes were further purified using iso-osmotic percoll solution. The purified monocytes were then incubated in RPMI supplemented with 5 % FBS for 30 min. Macrophages were differentiated from monocytes by incubation with granulocytes-macrophage colony stimulating factor (GM-CSF, 10 ng/ml, Bio Legend) in RPMI supplemented with 5% FBS for 5 days.

**Chemotaxis assay**

Migration of monocytes and macrophages against NDMV and NDTR were determined using μ-slide chamber (ibidi) following manufacturer’s instruction. The cell movement were visualized and captured by Nikon Eclipse Ni-U microscope using 20X objective. The movement of cells was then analyzed using ImageJ {Schneider:2012ui} and Chemotaxis/migration tool (ibidi).

**Polarization of differentiated THP-1**

THP-1 cells were purchased from Korean Cell Line Bank. THP-1 cells were maintained at 5 × 10^6^ cells/ml in RPMI (Gibco) supplemented with 10% heat-inactivated FBS (HyClone, GE healthcare Life Sciences) with 1% penicillin/streptomycin (Sigma-Aldrich). THP-1 cells were differentiated into M0 macrophages by treatment with PMA (100 ng/ml) for 48 h in RPMI supplemented with 10% FBS and 1% antibiotics. M0-differentiated THP-1 cells were exposed to NDTRs and NDMVs which were isolated from neutrophils (10^8^ cells) stimulated with fMLP (1 μM). For comparison, M0-differentiated THP-1 cells were stimulated with either LPS (100 ng/ml) and IFN-γ (10 ng/ml) for M1 polarization or IL-4 (10 ng/ml) and recombinant human IL-12 (10 ng/ml, Sinobiologicals) for M2 polarization. For evaluation of macrophage phenotype polarization, Polarized-THP-1 cells were detached by trypsin with EDTA (Life Technologies), fixed with Phosflow buffer (BD biosciences), and further stained with fluorescence-labeled anti-human monoclonal antibodies: CD14 (PerCP, BD biosciences), HLA-DR (APC, BD biosciences), CD80 (PE, Bio Legend), CD86 (PE, Bio Legend), CD209 (APC, Bio Legend), CD23 (APC, Bio Legend), CD163 (PE, Bio Legend), CD206 (FITC, BD biosciences), and CD1a (FITC, Bio Legend). Sample acquisition was performed using BD FACS caliber and was analyzed using FlowJo software. For evaluation of cytokine expressions, RNA was extracted from polarized-THP-1 cells using TRIzol (Ambion) and cDNA was synthesized using RT^2^ First strand kit according to manufacturer’s recommended protocols (Qiagen). RT-qPCR was performed by using cDNA (100 ng), primer (1 μl ) and RT^2^ SYBR Green qPCR mastermix (Qiagen) using Qiagen rotor. Briefly, The reactions were performed under the suggested cycling conditions: polymerase activation for 10 min at 95 ̊C, followed by 40 cycles of 15 s at 95 ̊C and 45 s at 60 ̊C. For all PCR experiments, post-PCR DNA melt curve analysis was performed to assess amplification specificity. DNA melting was carried out using a temperature ramping rate of 1°C per step with a 5-second rest at each step. Experiments were conducted in triplicate for reproducibility. Pre-designed RT^2^ qPCR primer assays (Qiagen) were used to determine the expression level of cytokines: iNOS (NOS2, PPH00173F), Arg-1 (PPH20977A), TNF-α (PPH00341F), TGF-β1 (PPH00508A), IL-10 (PPH00572C), IL-12A (PPH00544B), IL-6 (PPH00560C), and IL-1β (PPH00171C). Gene expression level were calculated as log_2_(2^-ΔΔct^) using GAPDH (PPH00150F) as a control .

**The uptake of neutrophil EVs by THP-1 cells**

Isolated neutrophil-derived EVs from cell tracker green (5 μg/ml, Invitrogen) stained neutrophils were exposed to M0-differentiated THP-1 cells on confocal dish (SPL Life sciences). Then live cell imaging was performed to visualize uptake of EVs by M0-polarized THP-1 cells using immunofluorescence microscopy at 37 ℃, 5% CO_2_ for 1 h. FACS acquisition was also performed to assess the uptake of neutrophil-derived EVs, M0-differentiated THP-1 cells were treated with isolated neutrophil-derived EVs for indicated time. The cells were harvested and fixed with phosflow fix buffer, permeabilized and stained with CD14 (PE) and CD66b (FITC). Sample acquisition was performed with BD FACS Calibur (BD Biosciences). The data was analyzed using FlowJo software (TreeStar Inc.).

**miRNA array and validation**

Total RNA was isolated from neutrophil EVs using the miRNeasy Micro Kit (Qiazen, USA) following the manufacturer’s protocol. The purity and concentration of the isolated RNA was measured with Nanodrop ND-1000 spectrophotometer (Nanodrop Technologies, USA). An equal amount (20 ng) of total RNA was converted to cDNA using the TaqMan MicroRNA Reverse Transcription Kit (Applied Biosystems, USA). 3 μl of RT product from total RNA was contained in a reaction volume of 10 μl with TaqMan Universial Master Mix II (Applied Biosystems) and TaqMan probe (hsa-miR-24-3p, hsa-miR-29a-3p, hsa-miR-122-5p, hsa-miR-126-3p, hsa-miR-150-5p, hsa-miR-451a, hsa-miR-1260a, hsa-miR-1285-5p, hsa-miR-4454, hsa-miR-7975) The expression levels of selected miRNAs were analyzed according to the TaqMan MicroRNA Assay protocol in triplicate. Cel-miR-39 was added in the samples during the RNA extraction step, and was used as an exogenous control for normalization of raw data. The miRNA profile of the samples was analyzed with Nanostring nCounter^TM^ system. Significantly, expressed miRNA (has-miR-1260a, hsa-miR-1285-5p, hsa-miR-4454, hsa-miR-7975, hsa-miR-126-3p, hsa-miR-150-5p, and hsa-miR-451a) were used to validate the effects of miRNAs on the phenotype polarization of macrophages. M0-differentiated THP-1(1 × 10^6^ cells) were transfected with 50 nM of indicated miRNA mimics (abm) using NEPA21 (NEPA gene) and incubated for 24 hrs in RPMI. RNA was extracted from transfected polarized-THP1 cells and the expression of iNOS and Arginase were determined by quantitative PCR as described earlier.

**Animal experiments**

Animal experiments were approved by the Institutional Animal Care and Use Committee of Kyungpook National University and Hallym University. BALB/c (male, 5-8 weeks old) mice were purchased from Orient Bio. Neutrophils were isolated from bone marrow by negative selection using either neutrophil isolation kit (Miltenyi Biotec) or EasySep^TM^ mouse neutrophil enrichment kit (StemCell technologies) as previously described {Park:2017gx}. EVs were isolated by stimulating mouse neutrophils (5 × 10^6^ cells/ml) with *E. coli* (1 × 10^6^ cells/ml). Isolated neutrophil derived EVs were dissolved in 100 μl phenol red-free RPMI and stored at -70 ℃. Experimental sepsis was induced in BALB/c mice by the cecal ligation and puncture (CLP) procedure as previously described {Yan:2004dx}. In brief, mice were anesthetized with pentobarbital (50 mg/kg, i.p.), and an abdominal mid-line incision was made. After laparotomy, the cecum was ligated below ileocecal valve, punctured with 22-gauge needle at two sides, and abdomen was closed. BALB/c mice were treated with either NDTRs (100 μl, i.p.), NDMVs (100 μl, i.p.) or vehicle (saline,100 μl, i.p.) 30 min before CLP surgery, and further treated on day 1, 2, and 3 after CLP surgery. The survival of septic mice were monitored for 9 days. DSS-induced chronic colitis was performed according to previous study {Marcon:2013iq}. In brief, BALB/c mice received water containing 2% DSS during first 5 days and further received normal drinking water for 10 days. At day 16, mice were exposed to water containing 2% DSS again until day 22. Mice were treated with NDTRs and NDMVs intraperitoneally on days 16, 18, 20, and 22. Mice were sacrificed on day 22 and colon size were measured, and the colons were prepared for histological analysis.

**Histological analysis**

For histological analysis, colons from mouse with DSS-induced colitis were fixed with 4% formaldehyde solution (Biosesang), and embedded in paraffin. Further, paraffin embedded samples were sectioned at 5μm-thickness and attached to adhesive glass slides (Matsunami), and stained with hematoxylin (Dako Mayer’s Hematoxylin, Agilent) and eosin (Daejung chemical and metal Co Ltd.). The HE stained tissues were visualized using Nikon Eclipse Ni microscope at 10X objective. Histological analysis were performed as previously described {Marcon:2013iq}. In brief, epithelial hyperplasia, mononuclear cells and polymorphonuclear cell infiltration in lamina propria, crypt inflammation, epithelial hyperplasia, and erythrocyte loss were scored.

**Serum evaluation**

Serum obtained from sepsis patients who admitted to the intensive care unit of Kyungpook National University Hospital between April and October 2016. Adults patients ( age ≥ 18) with sepsis in community-acquired pneumonia were enrolled. All patients provided informed consents in accordance with the Declaration of Helsinki. Venous blood was taken from septic patients within 24 h after ICU admission. Serum were obtained before neutrophil isolation and further stored at -70 ℃. Further, serum were thawed and filtered through 1.2 µM filter. The filtrate were then treated with 5µl of ExoQuick (System Biosciences), incubated for 30 mins at 4 ℃, followed by high-speed centrifugation (11,600 rpm for 1 hr). The pellets were then fixed with Phsoflow fix buffer and stained with Annexin V (FITC) and CD16 (PE) antibody. The sample were then acquired using BD FACS Calibur and the data analyzed using FlowJo.

**Statistical analysis**

Data are presented as the mean ± SEM for continuous variables and as the number (%) for the categorical variables. Data are compared using the student's t-test (unpaired, two-sided) and a p value of less than 0.05 was considered statistically significant. For the analysis of in vitro studies, statistical data were analyzed by Graphpad prism 7.0 (GraphPad Software Inc., San Diego, CA, USA). Comparisons between two groups were performed with either two-tailed Student's t test (parametric) or Mann-Whitney (non-parametric test). Survival data were analyzed using Mantel-Cox log-rank test. Values of P < 0.05 were considered to indicate statistical significance.

**References**
